# FoxP transcription factors are required for peripheral nervous system development

**DOI:** 10.64898/2026.09.22.753507

**Authors:** Hanna Yeliseyeva, Nikita Vladimirov, Esther T. Stoeckli

## Abstract

The FoxP family of transcription factors has been linked to neurodevelopmental disorders. FoxP1 and FoxP2 were identified as genes associated with autism spectrum disorders (ASD). FoxP1, FoxP2, and FoxP4 are expressed in many brain regions, including the cerebellum that has been of particular interest in the context of ASD. However, their role in the development of the peripheral nervous system (PNS) has been neglected. In view of the fact that a majority of ASD patients show some signs of sensory perception deficits that could go beyond the central nervous system, we focused on neural circuit formation in the PNS.

Our in vivo and in vitro studies revealed a role of all three FoxP transcription factors in neural circuit formation in the hindlimb of the chicken embryo. Silencing of FoxP genes altered motor axon guidance and also impacted neural crest cell migration. This in turn resulted in aberrant formation of dorsal root ganglia and deficits in neural circuits in the PNS.

Taken together, our results are in line with the hypothesis that some of the sensory perception deficits in ASD patients, either hyper-or hyposensitivity, are due to aberrant neural circuits in the PNS.

## Introduction

The FoxP proteins form a subfamily of the Forkhead box (Fox) transcription factors (Co et al., 2019; Lu et al., 2002; Shu et al., 2001; Teufel et al., 2003). Except for FoxP3, all family members, FoxP1, FoxP2, and FoxP4 are expressed in the nervous system during development. Their expression patterns are partially overlapping and highly dynamic (Ferland et al., 2003; Khouri-Farah et al., 2025; Takahashi et al., 2008). This adds to the complexity of FoxP function and the fact that we still know little about their role in neural circuit development. The FoxP transcription factors can regulate gene expression by forming homo-or hetero-dimers (Ahmed et al., 2024; Gao et al., 2023; Li et al., 2004a; Mendoza and Scharff, 2017; Sin et al., 2015). Their function is fine-tuned by various post-translational modifications (Gao et al., 2023). Furthermore, their activity and specificity in transcriptional regulation can be modified by specific binding partners (Estruch et al., 2018; Hickey et al., 2019). Often FoxP transcription factors act as repressors (Li et al., 2004a; Sin et al., 2015).

Based on clinical observations, FoxP1 haploinsufficiency has been linked to facial dysmorphology, intellectual disability and language impairment with or without autistic features (Meerschaut et al., 2017). Due to the variability of the phenotypes, the term FoxP1 syndrome has been generated (Siper et al., 2017, and references therein). FoxP2 has been identified as the cause of severe language disorders (Lai et al., 2001, 2003; Morison et al., 2023). The phenotypes seen in patients with either FoxP1 or FoxP2 mutations overlap, reflecting their interactions and the overlapping expression pattern in the developing brain (Bacon and Rappold, 2012). Less is known about the role of FoxP4 in human development. However, there are reports linking mutations in FoxP4 also to language disorders, as well as developmental delay and facial dysmorphologies (Del Viso et al., 2023; Snijders Blok et al., 2021).

Studies of the role of FoxP transcription factors in neural circuit development based on the analyses of human phenotypes have been focusing on FoxP2 mainly. A classical FoxP2 knockout mouse line revealed severe motor impairment, premature death by postnatal day 21 and absence of ultrasonic vocalization, a trait that is taken as replacement for language in mouse (Shu et al., 2005). Histological analysis of brain sections indicated cerebellar malformations. These were also found in heterozygous littermates, although to a lesser degree (French et al., 2007; Shu et al., 2005). These results were further supported by the analysis of a knock-in mouse that harbors the single amino acid exchange (R552H) that was identified in human language disorders (Fujita et al., 2008). Knockdown of FoxP2 by electroporation of shRNAs was shown to inhibit neurogenesis in the cortex, as transfected cells failed to migrate out of the ventricular zone (Tsui et al., 2013).

More recently, the function of FoxP1 in neural development has been addressed and revealed deficits in neural migration and differentiation as well. Knockdown of FoxP1 by in utero electroporation of short-hairpin RNAs (shRNAs) resulted in migration defects of cortical neurons and reduced dendritic branching (Li et al., 2015).

The function of Foxp4 has been studied in cerebellar slices, where knockdown of FoxP4 by shRNA resulted in reduced branching of Purkinje cell dendrites (Tam et al., 2011). Knockout mice die around E12.5 with severely impaired heart development (Li et al., 2004b).

The role of FoxP transcription factors in the wiring of the peripheral nervous system (PNS) is poorly understood. Two studies focused on the role of FoxP1 in motoneuron development (Dasen et al., 2008; Rousso et al., 2008). Changes in FoxP1 expression levels were linked to aberrant motoneuron differentiation and motor pool formation. In some nerves, aberrant connections or disturbed branching patterns in the forelimb were reported, while other nerves did not appear to be affected (Dasen et al., 2008). Similarly, FoxP2 and FoxP4 were shown to be required for motoneuron differentiation and detachment of motoneuron precursors from the apical surface (Rousso et al., 2012).

The PNS has been shown to contribute to altered sensory perception in human ASD patients (Tomchek and Dunn, 2007; Schaffler et al., 2019; Orefice, 2020) and mouse models for ASD (Orefice et al., 2016 and 2019). Because of their dynamic expression patterns that were only partially overlapping, we set out to analyze the role of all three FoxP transcription factors in PNS development. We used chicken embryos to silence FoxP1, FoxP2, or FoxP4 by in ovo RNAi followed by whole-mount staining, tissue clearing and light-sheet microscopy. Our in vivo studies demonstrated aberrant neural circuit formation in the PNS after silencing any one of the FoxP transcription factors. In agreement with their overlapping expression and their functional cooperation in the regulation of target genes, we found very similar phenotypes in hindlimb innervation and dorsal root ganglia formation. However, a more detailed analysis of the migration behavior of neural crest cells in vitro revealed some differences between the FoxP family members.

## Results

### FoxP genes are expressed in the developing neural tube of chicken embryos

To prepare for functional analyses of FoxP genes during neural circuit formation, we first studied their spatial and temporal expression patterns by in situ hybridization (ISH) on transverse spinal cord sections. We found distinct, but overlapping patterns of FoxP1, FoxP2, and FoxP4 during neural tube development, in line with previous reports (Dasen et al., 2008; Rousso et al., 2012). FoxP1 mRNA was detected in migrating neural crest cells (NCCs) and in cells lining the ventricles by HH18 (Fig. 1A,B). FoxP1 expression in motor neurons was very strong at HH24 (Fig. 1C,D). It was expressed also in some cells in the dorsal root ganglion although at lower levels than in motoneurons (Fig. 1C,D and C’,D’).

**Figure 1:**
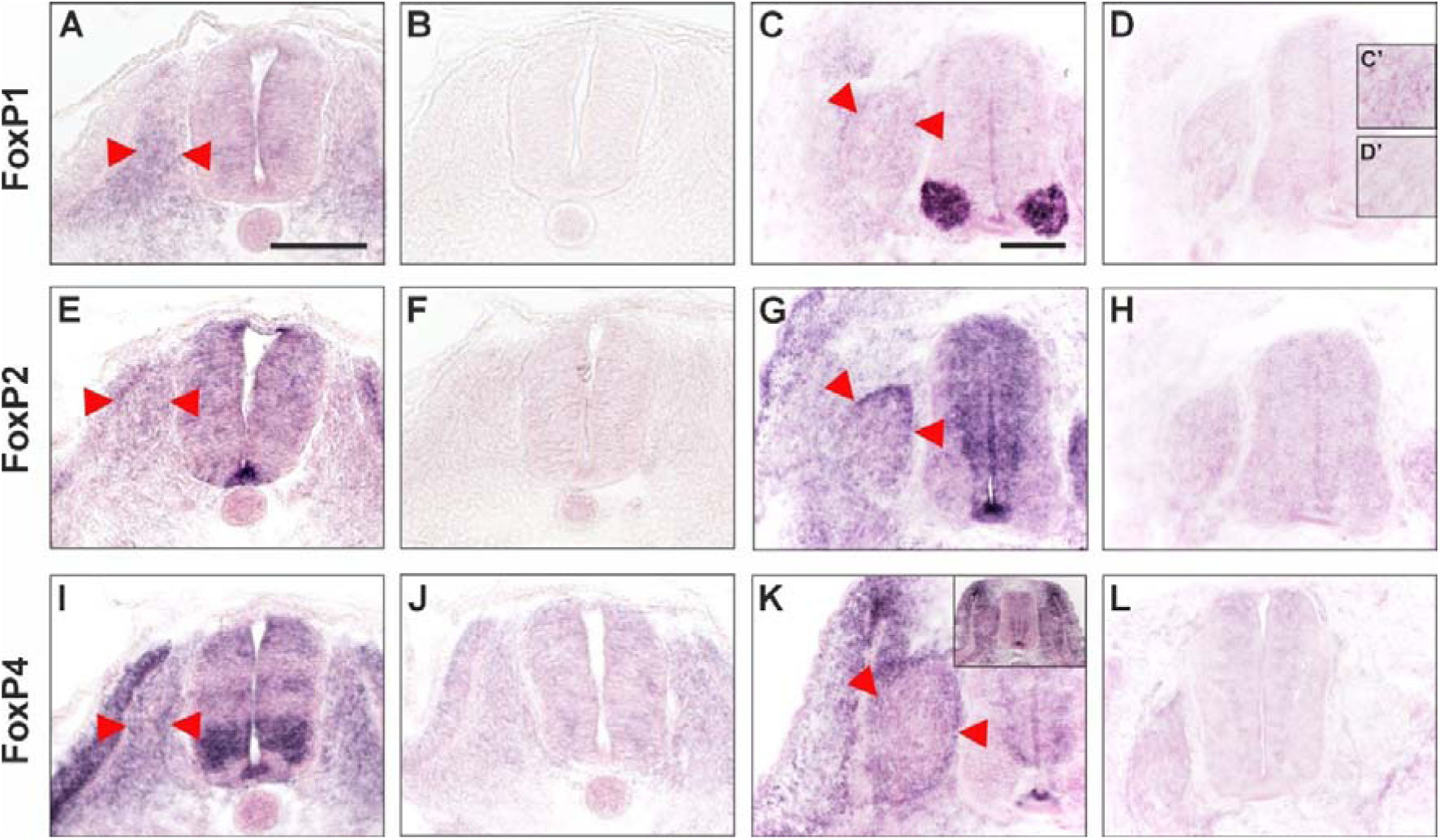
FoxP1, FoxP2 and FoxP4 mRNA expression in the lumbar chicken spinal cord. FoxP expression was analyzed in transverse spinal cord sections taken from the low thoracic and lumbar region and hybridized with antisense and sense RNA of FoxP1 (A-D), FoxP2 (E-H), or FoxP4 (I-L). FoxP1 expression was seen in migrating neural crest cells (NCCs) at HH18 (A; between red arrowheads). By HH24, the expression in motoneurons was very strong, but was also found in the dorsal root ganglia (DRG; red arrowheads in C and higher magnification in C’). FoxP2 was expressed strongly in the floorplate at HH18 (E). Expression was seen in migrating NCCs (between red arrowheads in E) and in cells throughout the neural tube. A relatively lower level of FoxP2 expression was maintained in some motoneurons at HH24 (G), but higher densities of FoxP2-expressing cells were clearly seen throughout the ventricular zone. In the DRG, expression was particularly strong in the dorso-medial area (red arrowheads in G). FoxP4 expression was strongest in motoneurons and motoneuron precursors at HH18, but also found in the floorplate and in a layer of interneuron precursors, as well as in the dorsal neural tube (I). Like the other family members, FoxP4 was expressed in migrating NCCs at HH18 (between red arrowheads). The expression pattern had changed considerably by HH24 (K). At this stage, FoxP4 was expressed in the floorplate, cells of the ventricular zone, and in cells throughout the DRG (red arrowheads in K). Very strong expression was found in cells dorsal to the DRG that were surrounding the dorsal end of the dermomyotome (see also insert in K). As controls, we used sense RNA of FoxP1 (B,D,D’), FoxP2 (F,H), and FoxP4 (J,L). Scale bars: 100 µm.

FoxP2 was expressed in the floorplate at HH18 and in neural precursor cells throughout the neural tube (Fig. 1E,F). FoxP2 mRNA was detected in migrating NCCs (between red arrowheads Fig. 1E) and later in the DRG, in particular in dorso-medial cells, at HH24 (Fig. 1G,H). At HH24, FoxP2 expression was also prominent in the ventricular zone but was weaker in mature neurons.

FoxP4 was dynamically expressed in the chicken spinal cord (Fig.1I-L). Its expression was observed in migrating NCCs at HH18 and in precursor cells of the neural tube. At HH18, these cells were concentrated in the dorsal area of the neural tube and in a more ventral layer. The strongest staining was seen in motoneuron precursors and in the floorplate (Fig. 1I,J). By HH24, FoxP4 expression was limited to the precursors in the ventricular zone and to the floorplate. Similar to FoxP1 and FoxP2, FoxP4 mRNA was also detected in the developing dorsal root ganglia (Fig 1K,L).

### Loss of FoxP1, FoxP2 and FoxP4 alters hindlimb innervation

Based on their temporal and spatial expression patterns, a role for FoxP family members in neural circuit formation was expected. We focused our functional analysis on the peripheral nervous system (PNS). Targeted downregulation of FoxPs was achieved using injection and electroporation of gene-specific dsRNA at E2 (see Experimental Procedures for details). Hindlimb innervation was assessed in cleared, 5-day-old, whole-mount chicken embryos stained with anti-neurofilament antibodies using a light-sheet microscope (mesoSPIM; Voigt et al., 2019; Vladimirov et al., 2024; Movies 1 and 2). Our subsequent analysis focused on morphologically well-defined hindlimb-innervating nerves, the ventral and dorsal crural nerves, the sciatic nerve, as well as the dorsal root ganglia.

To assess the efficiency of target gene silencing, we performed in situ hybridization (ISH) on spinal cord sections to quantify transcript levels (Fig. 2A). For FoxP1, for which an antibody was available, we also assessed protein levels by Western blots (Fig. 2B). We found a reduction of 34.9% at the mRNA level and a reduction of 39.1% at the protein level (Fig. 2A,B). For FoxP2 and FoxP4, we could only measure the reduction of the mRNA. We found a reduction of 39.3% for FoxP2 and 42.1% for FoxP4 (Fig. 2A). These values represent quite efficient downregulation. As we successfully transfect about 50% of the cells in the targeted area, these values suggest that our downregulation efficiency in the transfected cells is between 70% and 84%.

**Figure 2.**
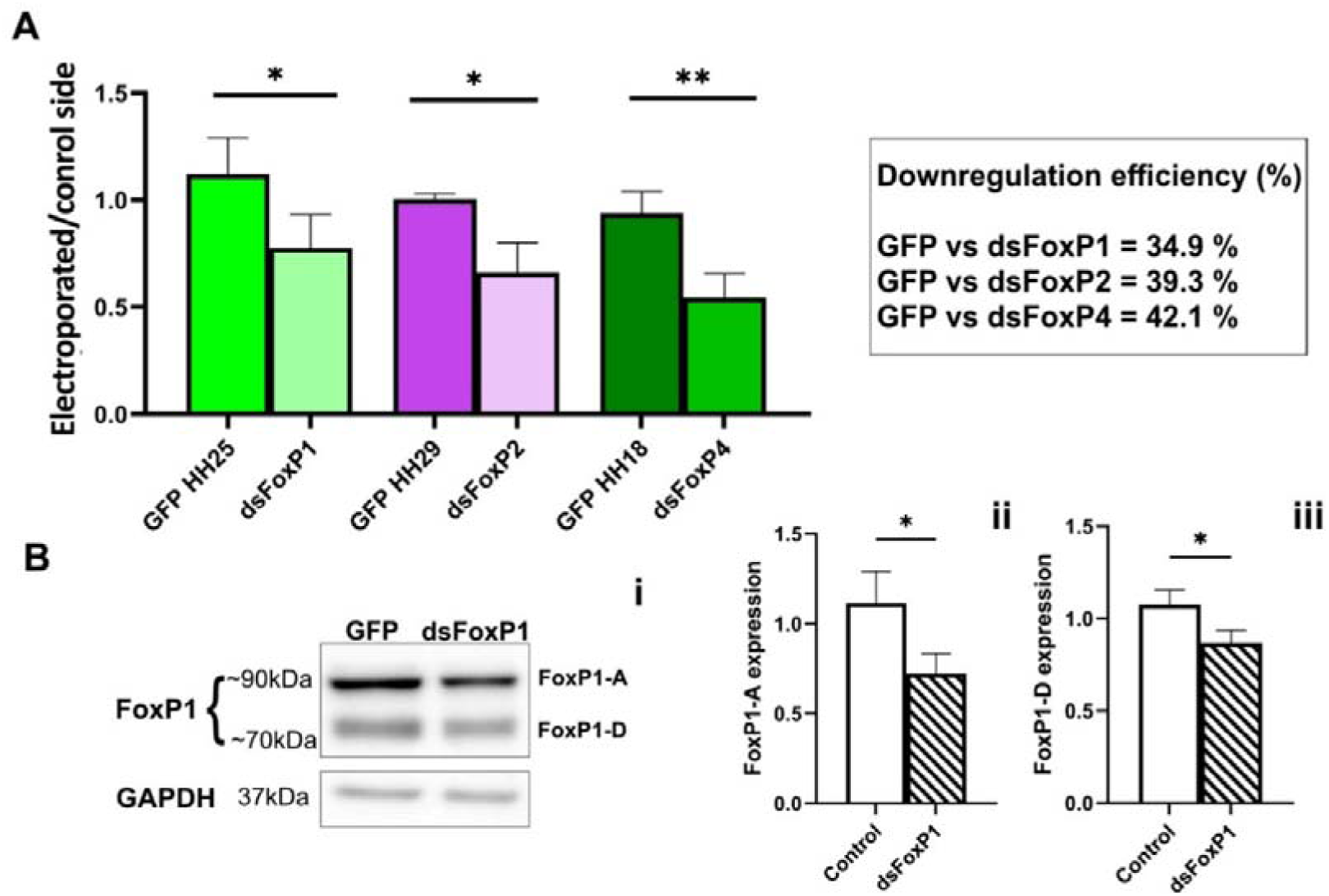
In ovo RNAi efficiently silences FoxP genes. (A) Downregulation efficiency was quantified by in situ hybridization after unilateral electroporation. Due to the dynamic expression patterns, different stages of embryonic development were used for the quantification of the specific FoxP expression levels. As expected, the ratio of the expression levels on the electroporated versus the non-electroporated side in control-treated, GFP-expressing embryos was always about 1. In contrast, the ratios of the expression levels on the electroporated compared to the non-electroporated control side were significantly lower for all dsFoxP embryos. We used n=5 GFP embryos compared to n=3 dsFoxP1 embryos at HH25; n=4 GFP embryos and n=3 dsFoxP2 embryos at HH29; n=4 GFP embryos and n=3 dsFoxP4 embryos at HH18. Welch’s t-test, **p < 0.01, *p < 0.05. (B) For FoxP1, we had an antibody to analyze the downregulation efficiency also at the protein level. The antibody stained two bands between 70 and 90 kD (Bi). The intensity of the upper band was decreased more strongly in embryos electroporated with dsFoxP1 (39.1%; Bii) compared to the lower band (20.5%; Biii). Three Western blots with different lysates were used for quantification. For each lysate, 3 GFP and 3 dsFoxP1 embryos, respectively, were used. Student T-test, *p < 0.05. (Graphpad Prism).

The quantitative analysis of the sensory and motor components of the PNS in the hindlimb of whole-mount embryos imaged with mesoSPIM consistently showed aberrant innervation patterns after silencing FoxP genes (Fig. 3). Specifically, the sciatic nerve was strongly affected in experimental embryos compared to control embryos, both wildtype and embryos injected and electroporated only with the GFP-expressing plasmid. In experimental embryos lacking either FoxP1, FoxP2, or FoxP4, the sciatic nerve failed to segregate properly into an anterior and a posterior branch at a ratio of about 1:1 as seen in controls (Fig. 4A,B). In embryos lacking FoxP expression, some axons corrected their initial decision (orange arrowheads in Fig. 4C,D). In addition, after silencing FoxP genes, we often found duplications in the origin of a branchlet that extended dorsally along the posterior branch of the sciatic nerve (white arrowheads in Fig. 4A-E). We compared the embryos lacking FoxPs to control-treated embryos that were injected and electroporated only with the GFP-expressing plasmid (GFP controls). This plasmid was used as a transfection control in all our treated embryos. The GFP controls were not different from the non-treated control embryos, which were also included in our analysis to rule out effects from our experimental procedures. For quantification, we compared the percentage of embryos per group that did not show any deficits in the extension of the sciatic nerves (Fig. 4F,G).

**Figure 3.**
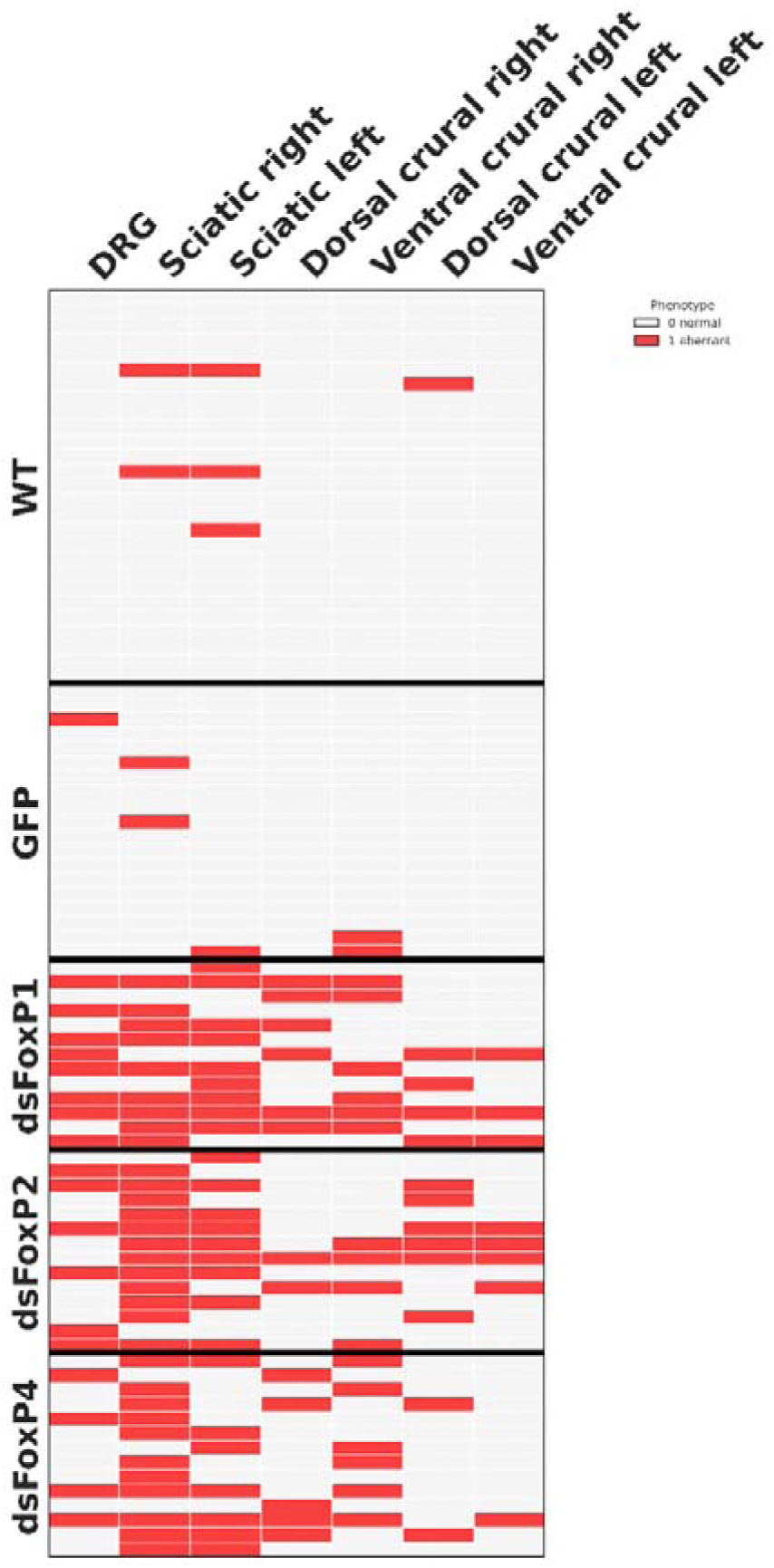
Quantitative analysis of FoxP loss-of-function phenotypes. We analyzed n=27 untreated embryos, compared to control-treated, GFP-expressing embryos (n=19) and dsFoxP1 (n=13), dsFoxP2 (n=14), as well as dsFoxP4 embryos (n=14). Each line represents one embryo. The analyzed structure is given on top. Normal phenotypes are represented in grey, aberrant phenotypes are represented in red.

**Figure 4.**
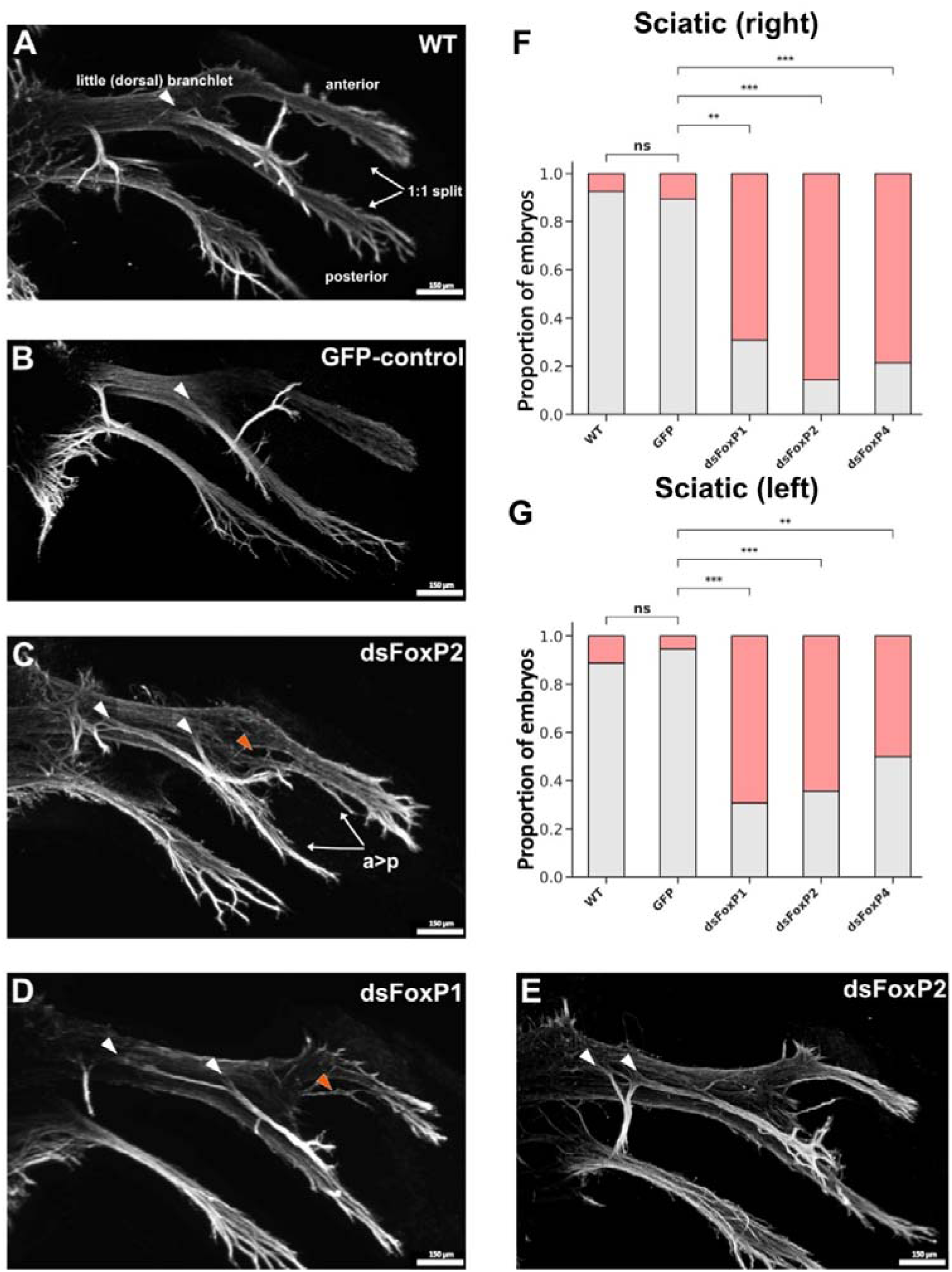
Loss of FoxP transcription factors leads to deficits in the development of the sciatic nerve. (A) Overview of the normal morphology of the dorsal part of the sciatic nerve in a HH25 WT embryo. Axons are segregated equally into an anterior and a posterior branch. A little dorsal branchlet extends alongside the posterior branch and has a single origin (white arrowhead). (B) In an age-matched embryo electroporated with only the GFP expression plasmid, the morphology of the sciatic nerve is preserved. (C-E) In contrast, embryos lacking either one of the FoxP transcription factors showed aberrant development of the sciatic nerve. Because the phenotypes did not differ between the experimental groups, we show selected examples. Consistently axons were unequally segregated into the anterior and posterior branches (C). Often this was observed in combination with axons correcting their choice and therefore changing from the posterior to the anterior branch (orange arrowhead in C and D). Very common was also the aberrant formation of the little dorsal branchlet, as part of it split off the main sciatic trunk too early, at a proximal position. This resulted in a duplication of the origin in experimental compared to control embryos (white arrowheads in A-E). Scale bars: 150 µm. mesoSPIM imaging, with Napari rendering. (F,G) Silencing FoxP genes resulted in a reduction of correctly innervated limbs. The proportion of embryos with normal morphology is given as a grey bar for the right side (F) and the left side (G). We used n=19 GFP embryos, n=13 dsFoxP1, n=14 dsFoxP2, and n=14 dsFoxP4 embryos. Fisher test, ***p < 0.001, **p < 0.01. Python.

We also consistently found aberrant branching of the crural nerves in the absence of FoxP transcription factors (Fig. 5). In untreated control embryos (Fig. 5A) and in GFP controls (Fig. 5B), the ventral crural nerve formed a smooth bundle that was connected to the dorsal nerve with a thin axon bundle. After silencing of FoxP1 and FoxP4, the ventral crural nerve was defasciculated to various degrees (blue arrowheads, Fig. 5D). The characteristic branching pattern of the dorsal crural nerve (Fig. 5A,B) was strongly reduced in embryos lacking FoxP transcription factors (orange arrowheads; Fig. 5C-E). In some cases, an overall retardation of hindlimb innervation was found, as seen in Fig. 5E, where both the dorsal and the ventral crural nerve were much thinner and delayed in their development. Silencing FoxP2 did not induce significant changes in the crural nerves.

**Figure 5.**
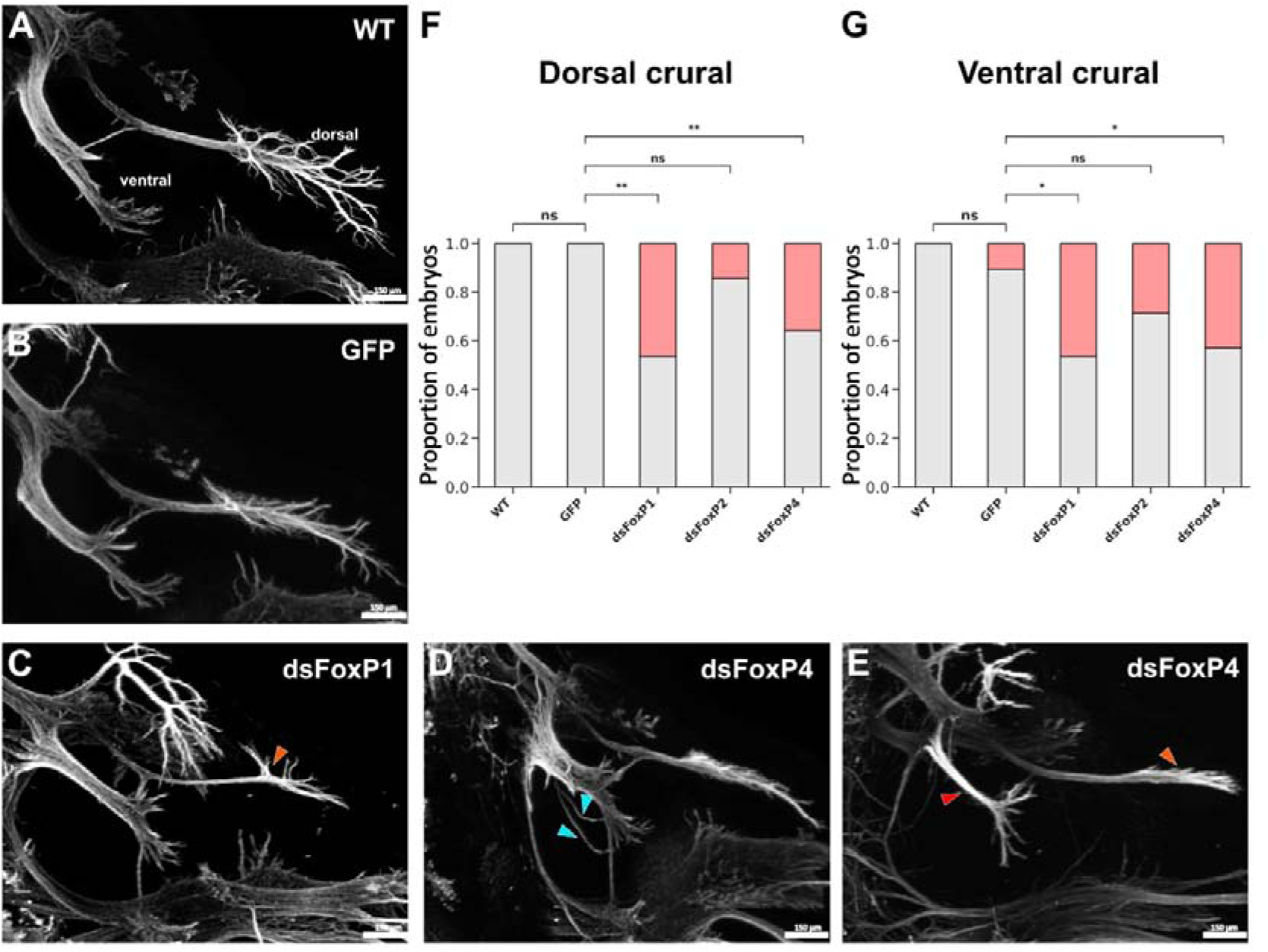
FoxP silencing interferes with the normal morphology of ventral and dorsal crural nerves. (A) At HH25, the ventral and dorsal crural nerves have extended into the hindlimb in wildtype embryos. (B) No difference was seen in embryos injected and electroporated with the GFP expression plasmid alone. (C-E) Silencing of either FoxP1 or FoxP4 induced aberrant innervation patterns of the crural nerves. Very common were aberrant branching of the dorsal crural nerve (orange arrowhead in C and E). Instead of forming a smooth ventral nerve, downregulation of FoxP1 or FoxP4 induced defasciculation (blue arrowheads in D). In some embryos after silencing FoxP genes, we observed a delay in hindlimb innervation, resulting in thin branches (red arrowhead) and absence of age-appropriate branching (orange arrowhead in E). Scale bars: 150 µm; mesoSPIM imaging, with Napari rendering. (F,G) Silencing FoxP genes resulted in a reduction of correctly formed crural nerves. The proportion of embryos with normal morphology is given as grey bar for the right side (F) and the left side (G). We used n=19 GFP embryos, n=13 dsFoxP1, n=14 dsFoxP2, and n=14 dsFoxP4 embryos. Fisher test, **p < 0.01, *p < 0.05. Python.

Aberrant forelimb innervation in the absence of FoxP1 was described previously (Dasen et al., 2008). The observed reductions in branching of peripheral nerves are thus in agreement with our findings of reduced branching in the hindlimb nerves.

### Loss of FoxPs changes DRG morphology

Strikingly, we found aberrant phenotypes in both left and right legs despite unilateral electroporation (Fig. 3 - 5). This can be explained by a contribution of sensory axons to both the crural and the sciatic nerves. Because of the timing of our electroporation, we also target neural crest cells before delamination. Delaminating NCC migrate to both sides, therefore, they also contribute to a loss-of-function phenotype on the non-electroporated side.

Indeed, when we analyzed the formation of dorsal root ganglia, we found evidence for a problem with neural crest cell migration. While there was no difference between the untreated and the GFP control group, we found aberrant alignment of the DRG and a marked reduction and variability in size after downregulation of FoxP transcription factors (Fig. 6). Often, adjacent DRG were merged or formed joint roots (asterisks, Fig. 6Aiv). Our quantification of DRG morphology and alignment revealed one out of ten embryos of the GFP control group with a problem in DRG formation. None of the non-treated embryos had any problems with DRG morphology or alignment. In contrast, after silencing FoxP1, we found only 33.3% (4 out of 12) of the embryos with normal DRG. After silencing FoxP2, 58.3% of the embryos had normal DRG (7 out of 12), a value that was very similar in embryos lacking FoxP4, where we found normal DRG in 55.6% (5 out of 9) of the embryos at age HH24-25. By utilizing Axonin1/Contactin-2 staining together with a boundary cap cell (BCC) marker, we were able to detect changes in dorsal BCC cluster arrangement along the neural tube (Fig. 6A). All the observed changes are in line with aberrant NCC migration and clustering. We found marked reduction in DRG size and a reduced number of roots extending towards the spinal cord also in embryos stained with antibodies against Neurofilament (Fig. 6B). Strikingly, we found a ‘zig-zag’ pattern of DRG along the spinal cord due to the non-symmetrical arrangement along the anterior-posterior axis.

**Figure 6.**
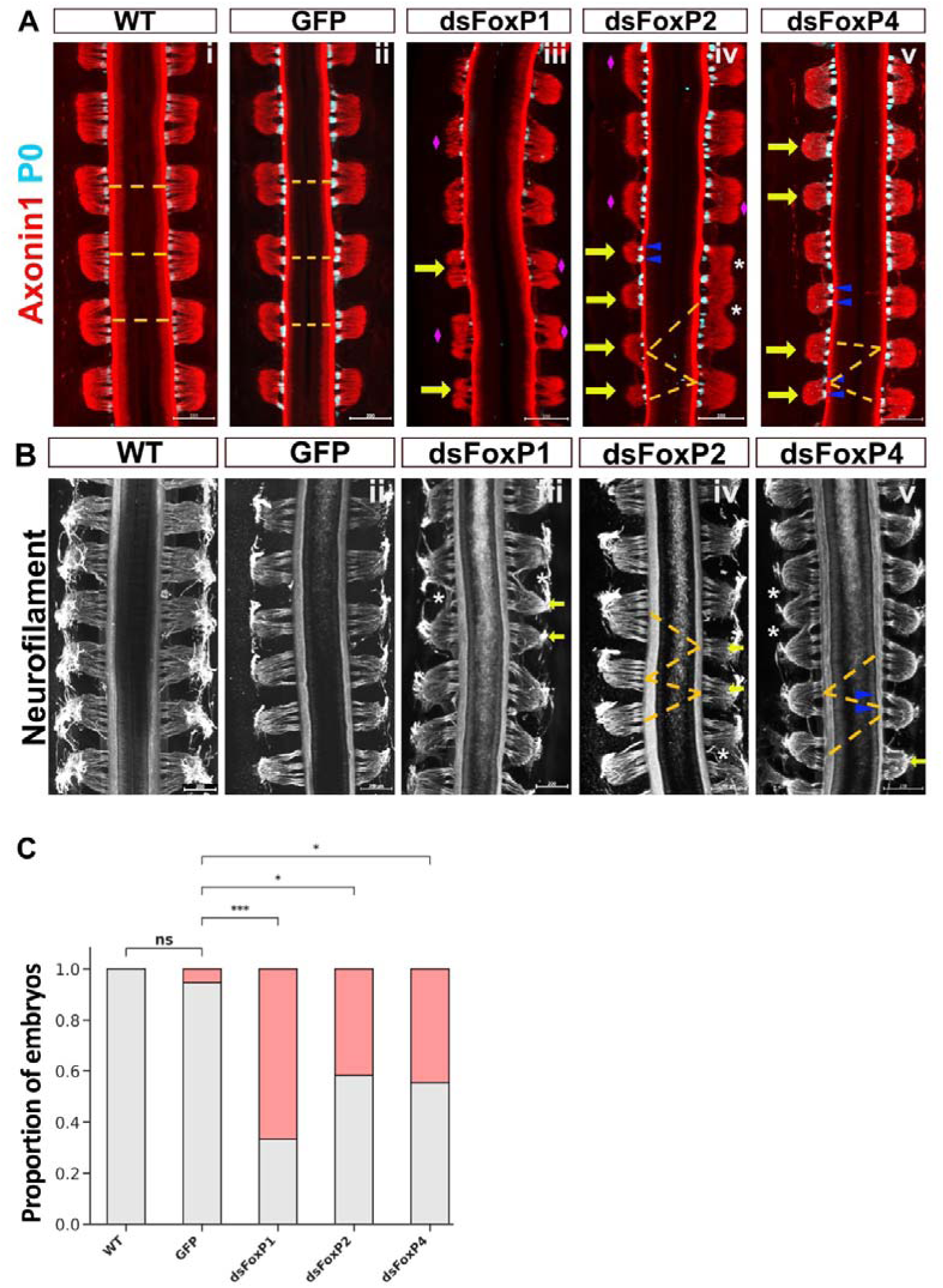
FoxP silencing leads to the aberrant formation of DRG. (A) Whole-mount embryos sacrificed at HH24 and stained for Axonin-1/Contactin-2. Boundary cap cell clusters were stained with an antibody against P0 to highlight the entry of dorsal roots into the spinal cord. In wildtype (WT; i) and in GFP control embryos (ii), dorsal root ganglia were arranged symmetrically along the neural tube (indicated by yellow dashed lines). Their sizes were comparable, and they were all sending between 4 to 6 roots towards the neural tube. In contrast, after silencing FoxP1 (iii), FoxP2 (iv), or FoxP4 (v), the variability in DRG sizes was striking. Their arrangement was completely messed up (yellow dashed lines). Adjacent DRGs sometimes merged into one big structure (asterisks). The number of roots per ganglion was often markedly reduced (blue arrowheads). The shape of the ganglia deviated from the bell shape that is found in controls. (B) The same phenotypes were also seen after staining with anti-Neurofilament antibodies. Also here, the arrangement of the DRG was often a zig-zag pattern (yellow dashed lines), DRG sizes varied, and the number of roots was often reduced. Scale bar: 200Iµm. mesoSPIM imaging, with Napari rendering. (C) Downregulation of FoxP genes resulted in aberrant DRG phenotypes in all experimental groups. Fisher test, ***p < 0.001, *p < 0.05. Python.

### Silencing of FoxP genes affects neural crest cell delamination and migration

Based on our findings of aberrant size and positioning of DRG, we focused on NCC migration. To follow NCCs in vivo, we performed whole-mount staining for HNK1, a marker for NCCs, at an earlier developmental stage (HH18). Using HNK1 staining together with ECi-based tissue clearing, we were able to assess the delamination pattern of NCCs as well as their polarity in whole-mount embryos. Silencing FoxP genes affected polarity and migration in all experimental groups (Fig. 7). As NCC delaminate from the neural tube, they undergo epithelial-to-mesenchymal transition and change shape from a circular to an elongated morphology (Theveneau and Mayor, 2012; Giovannone et al., 2015). The analysis of young whole-mount embryos suggested migration deficits in delaminating neural crest cells, in agreement with the observed DRG phenotypes at later stages of development.

**Figure 7.**
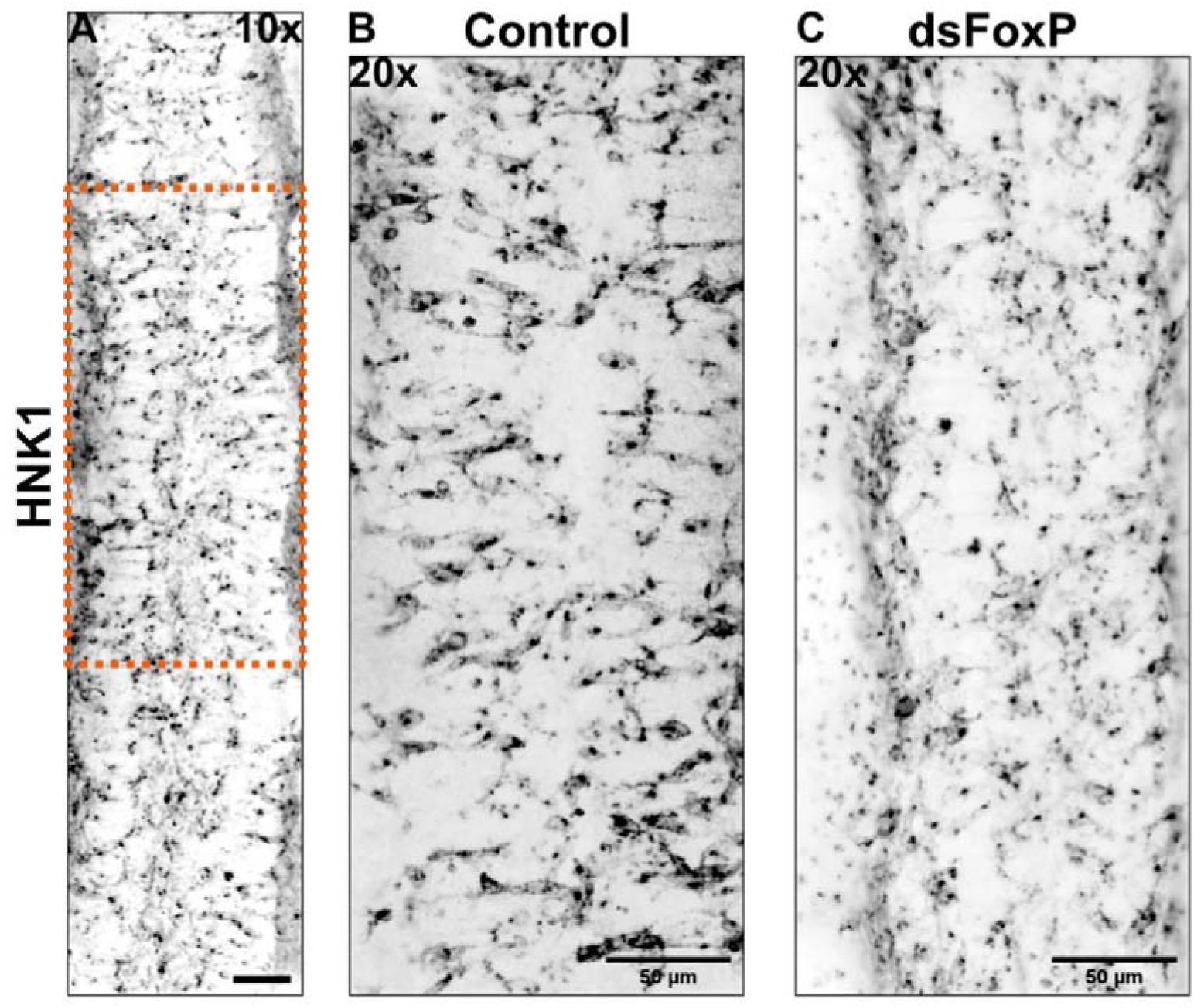
The loss of FoxP genes interferes with neural crest cell delamination and migration. (A-B) Migrating NCCs stained with an anti-HNK1 antibody in a HH17 whole-mount embryo at low (A) and higher magnification (B). NCCs are polarized and migrate perpendicular to the dorsal midline towards either side of the spinal cord. (C) In contrast, in the absence of FoxP transcription factors, NCCs are not polarized and fail to migrate efficiently to the sites where DRG form. Scale bar: 50 µm. Fiji/ImageJ.

To assess the dynamics of NCC migration upon epithelial-to-mesenchymal transition in detail, we used in vitro assays. For this purpose, we cultured neural tube explants taken from the low thoracic-lumbar region of HH17 embryos and followed NCC migration with time-lapse imaging. NCCs were visualized by tdTomato that was expressed due to injection and electroporation of crestin::tdTomatoF at HH12. We assessed the migratory behavior of NCCs during the first 20h after delamination. Live imaging of NCCs was started 2 hours after plating. In cultures of explants taken from control-treated (GFP) embryos, cells were very motile and switched between an elongated and a more roundish morphology. As described before (Carmona-Fontaine et al., 2008; Kasemeier-Kulesa et al., 2005; Li et al., 2019), NCCs were touching each other via transient processes and moved in different directions after contact. Using manual tracking, we followed the movements of selected NCCs and reconstructed their pathways using custom Python coding (Fig. 8). We quantified the total distance travelled (sum of all segments: Fig. 8A), as well as the absolute displacement of the cells (distance measured between location at t=14h and t=20h; Fig. 8B). This allowed us to calculate the straightness index (absolute displacement divided by total distance travelled; Fig. 8C) and the average migratory speed (Fig. 8D).

**Figure 8.**
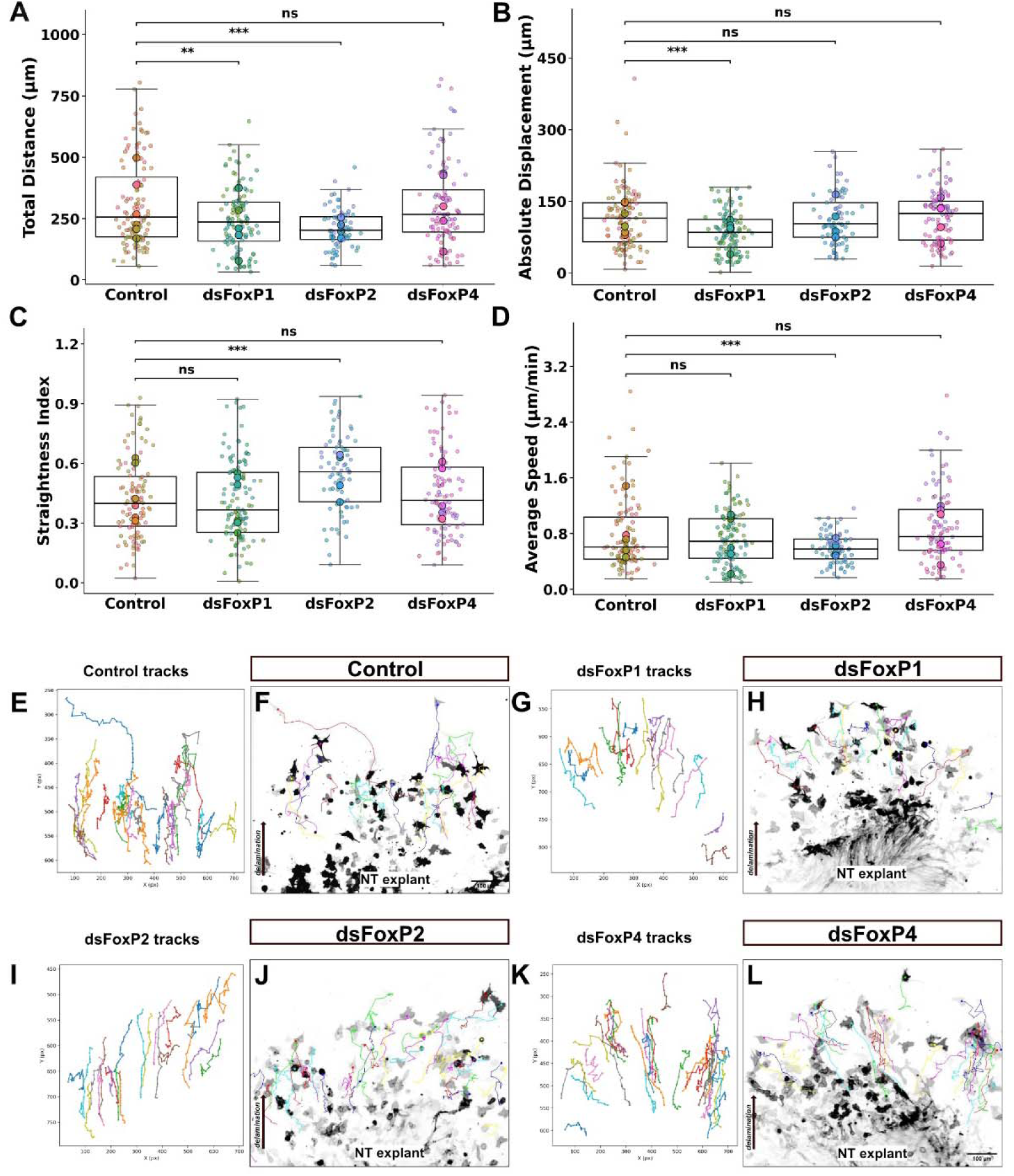
FoxP silencing affects lead NCC behavior upon EMT in a gene-dependent manner. (A) The total distance travelled was measured by adding the distances covered between frames. Small dots indicate measurements of individual cells, bigger dots represent the average per explant. All measurements belonging to the same explant are represented by the same color. (B) Absolute displacement refers to the length of the straight line drawn from the position of the cell in frame 1 to the position of the cell at the end of the observation period. (C) The straightness index is calculated by dividing the absolute displacement by the total distance travelled. A value of 1 would be migration on a straight line, values below 1 indicated that cells moved back and forth and deviated considerably from a straight line. (D) The average speed is calculated as the total distance traveled divided by the time of live imaging. (E,F) Example of cells migrating away from an explant taken from a control embryo (E, tracks only, F, tracks overlayed with frame 1 of the respective movie). (G,H) Example of tracks observed for NCC from explant taken from a dsFoxP1 embryo. (I,J) Tracks seen for NCCs from an explant taken from a dsFoxP2 embryo. (K,L) Tracks seen for NCCs migrating from an explant taken from a dsFoxP4 embryo. Welsch T-test ***p < 0.001, **p < 0.01, *p < 0.05. Kaggle, Python. Fiji, manual tracking. Scale bars: 100 µm.

Downregulation of FoxP1 consistently impaired neural crest cell motility relative to controls. Both the absolute displacement and the total distance covered were reduced (Fig. 8A,B, Movie 3), but the straightness index and the average speed were unchanged (Fig. 8C,D). In contrast, NCCs lacking FoxP2 had the same displacement as controls, but the total distance that they covered was shorter than for controls, resulting in slower speed (Fig. 8D) but increased straightness index (Fig. 8C). No difference to control cells was seen for those lacking FoxP4. The changes in migration behavior can be appreciated when comparing traces of cells taken from control explants (Fig. 8E,F; Movie 4) with cells from explants lacking one of the FoxPs (Fig. 8G-L; Movies 5-7). In control cultures, cells interacted with other cells via rapidly extending processes (Fig. 8F). Cells changed direction after bumping into each other and changed morphology. A similar behavior, but shorter traces were seen after silencing FoxP1 (Fig. 8G,H). Cells lacking FoxP2 reached the same endpoints but covered less distance to get there, explaining the straighter traces (Fig. 8I,J). No difference in displacement or total distance traveled compared to control cells was seen after silencing FoxP4 (Fig. 8K,L).

In control cultures, NCCs contacted each other via transient processes and rapidly split again to migrate in different directions, as described previously (contact inhibition of locomotion; Stramer and Mayor, 2017; Fig. 9A). A different behavior was seen for cells lacking FoxP2 (Fig. 9B,D). The cells were more roundish than control cells. They were motile and reached the same displacement values as controls, although the total distance was shorter, and as a consequence, the straightness index was higher than for controls. This is reflected in fewer cell-cell contacts (Fig. 9B). Yet another behavior was seen for cells lacking FoxP4. They contacted other cells not via rapidly extending processes, but stayed in contact with other cells by forming tight contacts over large surface areas (Fig. 9C,D). This clustering behavior was not interfering with migration as the displacement and the total distance were not different from controls, despite the fact that the cells moved as a cluster.

**Figure 9.**
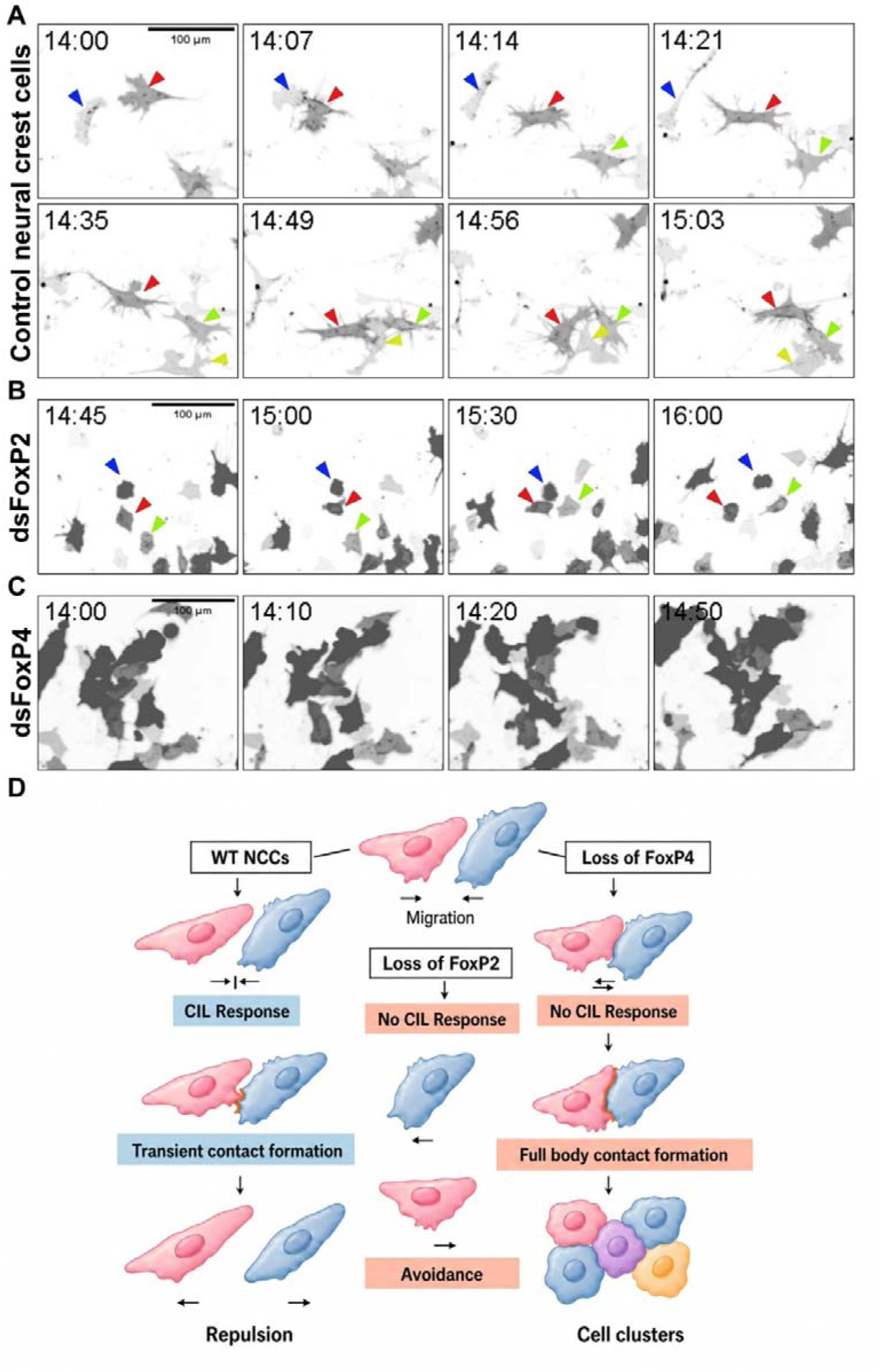
Silencing FoxP genes results in abnormal locomotion behavior of NCCs. (A) Control NCCs extend processes to contact other NCCs. After contact, cells separate and move in different directions. Contacts are often repeated between the same cells (cells are labeled with the same color throughout the image series, representing 63 minutes of live imaging). (B) In the absence of FoxP2, cells are also motile, despite the more roundish morphology. They move with fewer contacts, reflected by a higher straightness index (see Figure 8A-C). (C) Cells lacking FoxP4 do not differ from control cells with respect to displacement and total distance covered. However, their behavior observed by live imaging for the same time period clearly differs the most. Rather than initiating transient contacts via processes, they tend to stick together via extended cell-cell contacts involving a large surface area. (D) Schematic summary of the observed phenotypes. Control cells make contacts via processes in a transient manner. Along their path, they contact each other repeatedly. Loss of FoxP2 reduces the likelihood of cell-cell contacts. Cells are more roundish and produce fewer processes. In the absence of FoxP4, cells do not contact each other via transient processes, but rather form tight cell-cell contacts by including large parts of the cell surface. Scale bars: 100 µm.

Overall, the data suggest that all FoxPs are involved in the regulation of NCC migration. However, the mechanisms underlying the changes are different. While FoxP1 loss of function induced the least difference in behavior compared to controls, loss of either FoxP2 or FoxP4 changed the behavior strongly. Interestingly, the quantitative difference between controls and dsFoxP4 cells was not significant, despite the fact that the morphology of the cells and the contact behavior were very different (see Fig. 9D). However, the clustering made the quantitative assessment of the behavior of single cells more difficult.

### FoxP1, FoxP2 and FoxP4 re-expression in the developing neural tube rescues PNS phenotypes

In order to confirm the specific role of FoxP genes in the establishment of peripheral neural circuitry, we sought to rescue the aberrant phenotypes obtained after FoxP1, FoxP2 and FoxP4 downregulation by injecting a plasmid containing the full-length mouse FoxP1, FoxP2, or FoxP4 cDNA, respectively (Fig. 10). Because FoxP transcription factors can form homo-or heterodimers, we expected that both too little and too much of any one of them would have detrimental effects on hindlimb innervation and DRG development rather than restoring the normal phenotype. Therefore, we tested different concentrations of the rescue constructs. E2 chicken embryos were injected and electroporated unilaterally with 20 ng/µl β-actin::GFP, 500 ng/µl of dsRNA against one of the FoxP genes and 50, 100, or 250 ng/µl of the respective rescue plasmid driven by the β-Actin promoter. The expression of the β-actin::mFoxP1-3xHA, β-actin::mFoxP2-3xFLAG, and β-actin::mFoxP4-6xmyc constructs in spinal cord neurons and DRG was confirmed by immunostainings of transverse spinal cord sections of embryos sacrificed at HH25 (Supplementary Figure 1).

**Figure 10.**
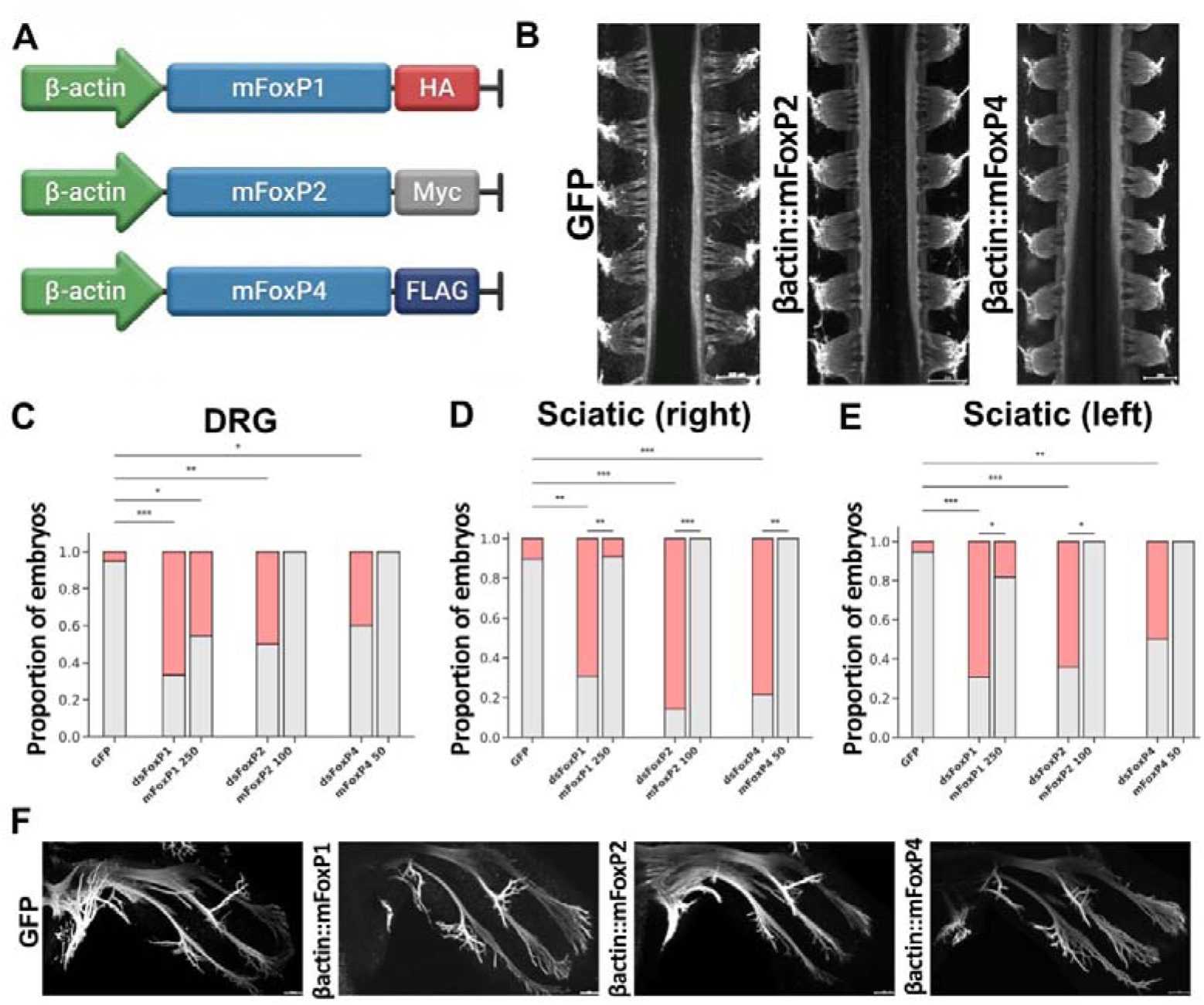
The phenotypes observed in the absence of FoxP proteins can be rescued by expression of the respective mouse cDNAs. (A) As rescue constructs, we used the respective mouse cDNAs tagged with either HA (FoxP1), myc (FoxP2), or Flag (FoxP4). All constructs were expressed under a β-actin promoter. Because FoxP transcription factors can form homo-and heterodimers, the rescue was dose-dependent. Too much or too little expression of any one of the FoxPs was expected to have a detrimental effect. (B) Examples of DRG images. mFoxP2 and mFoxP4 expression rescued DRG morphology in dsFoxP2 and dsFoxP4 embryos, respectively. They were not different from GFP controls. mesoSPIM imaging, Napari rendering; (C) Low to intermediate levels of FoxP2 and FoxP4 re-expression were sufficient to restore normal morphology of DRG. The phenotype induced by dsFoxP1 could not be rescued by expression of even a high dose of the mFoxP1 construct; (D-E) Low to intermediate levels of FoxP1, FoxP2 and FoxP4 re-expression were sufficient to restore normal morphology of sciatic nerves. The right (D) and the left side (E) were quantified separately also for the rescue experiments. (F) Examples of successfully rescued sciatic nerves. The morphology after rescue was not distinguishable from GFP controls. Mesospim imaging, Napari rendering. Hindlimb N: n=13 dsFoxP1, n=14 dsFoxP2, n=14 dsFoxP4, n=19 GFP, 50. DRG N: n=12 dsFoxP1, n=13 dsFoxP2, n=9 dsFoxP4, Rescue N: n=11 β-actin::mFoxP1 250 ng/µl, n=6 β-actin::mFoxP2 100 ng/µl, n= 6 β-actin::mFoxP4 ng/µl.

As reported above, silencing of all FoxPs markedly disrupted PNS formation. When we co-injected the rescue constructs β-actin::mFoxP2-3xFLAG (100 ng/µl) and β-actin::mFoxP4-6xmyc (50 ng/µl), we could fully restore normal DRG morphology in embryos treated with dsFoxP2 or dsFoxP4, respectively (Fig. 10B,C). In contrast, none of the concentrations of the FoxP1 rescue plasmid (β-actin::mFoxP1-3xHA) that we tested was able to restore the normal DRG phenotype (Fig. 10C). We hypothesize that the incomplete rescue of aberrant DRG phenotypes, at any of the tested concentrations, can be explained by the known requirement of FoxP transcription factors to act as dimers. High doses of the rescue plasmid may disrupt normal heterodimer formation and thus change the normal gene expression program. Notably, not all cell types require the same FoxP-dependent transcriptional programs. All tested concentrations of the rescue plasmids were effectively rescuing the aberrant phenotype in the sciatic nerve (Fig. 10D-F). For FoxP4 even the lowest concentration of 50 ng/µl of the rescue plasmid was effective (Fig. 10D,E).

Taken together, our studies support a role of FoxP transcription factors in PNS development. Our in vitro and in vivo studies indicate a collaborative function of FoxP1, FoxP2, and FoxP4 in hindlimb innervation. In contrast to earlier studies focusing on motoneurons, we expand our understanding of the role of FoxP transcription factors to neural crest cell development. Future studies will need to analyze later stages of PNS circuit development to reveal the specific effects of the different FoxP family members.

## Discussion

In this study, we demonstrate that FoxP transcription factors play a role in the formation of neural circuits in the peripheral nervous system (PNS). Using temporally controlled in ovo RNAi in the chicken embryo combined with whole-mount tissue clearing and light-sheet mesoSPIM microscopy, we show that silencing FoxP1, FoxP2, or FoxP4 leads to reproducible and nerve-specific defects in axon guidance and neural crest cell migration, which is required for DRG organization. As we analyzed only young stages of PNS circuit development, the effects of silencing the different FoxP family members did not result in gene-specific phenotypes yet. Or, in other words, the specific effects of FoxP loss of function were not detectable with the depth of our analysis. An exception is the detailed analysis of neural crest cell migration behavior. Here, we clearly detected differences between the experimental groups. However, the aberrant migration behavior did not manifest in specific differences of DRG formation and position between the experimental groups. DRG formation was always affected compared to controls, no matter how the migratory behavior was changed.

Our analysis of hindlimb innervation revealed that FoxP downregulation disrupts the morphology of major peripheral nerves, particularly the sciatic and crural nerves. Peripheral nerves are “mixed” structures comprising axons of NCC-derived sensory neurons and glia, as well as CNS-derived motor axons. The observed phenotypes, including branching defects, axonal misrouting with correction attempts, and defasciculation, suggest that FoxP transcription factors control the expression of proteins that are required for axon guidance decisions at key choice points. These findings are consistent with previous reports implicating FoxP1 in motor neuron connectivity and target selection (Dasen et al., 2008; Palmesino et al., 2010). FoxP2 and FoxP4 have been implicated in motoneuron differentiation (Rousso et al., 2012). Here, we extend these observations to study the role of FoxP1, FoxP2, and FoxP4 in PNS development. In particular, the contribution of FoxP transcription factors to the sensory part of neural circuits in the PNS has not been appreciated before.

While our observations for loss of FoxP1 were consistent with the motor axon deficits described previously (Dasen et al., 2008), the fact that we saw aberrant hindlimb innervation on both sides after unilateral electroporation could only be explained by a contribution of NCCs and their derivatives to the phenotype. After delamination from the neural tube, NCCs choose either side. Therefore, we analyzed younger embryos to assess the formation of DRG. Trunk neural crest cells migrate in segmental waves through the anterior part of somites along the embryonic axis (Giovannone et al., 2015; Teddy and Kulesa, 2004; Theveneau and Mayor, 2012). They move as organized chains composed of leader and follower cells and ultimately give rise to peripheral neurons (including dorsal root ganglia and sympathetic neurons) and glial cells, the Schwann cells. Their eventual fate is closely tied to their migratory path and position, shaped by somite boundaries and reinforced by intrinsic differences in cell identity (Richardson et al., 2016).

Indeed, a role of FoxP transcription factors in NCC migration was supported by the aberrant arrangement and the variability in size of DRG in experimental compared to control embryos (Fig. 6). Although the DRG phenotype looked similar for all dsFoxP embryos, the detailed analysis of their effects on NCC migration revealed specific traits that were altered (Fig. 9). In the absence of FoxP1, NCCs migrated less far, but their migratory behavior was similar to control cells. In contrast, in the absence of FoxP2, NCCs moved less but straighter than control cells. This was accompanied with a difference in contact behavior (Fig. 9). In contrast to wildtype cells, which contacted each other repeatedly via transient processes, FoxP2-deficient NCCs were roundish and failed to make the typical processes to induce contacts with adjacent cells. Finally, NCCs lacking FoxP4 revealed the opposite behavior rather than inducing recurrent and transient contacts, they stuck together after contact and maintained cell-cell contacts involving a large proportion of their surface. As a consequence, they moved together as a group rather than individual cells. This made the quantifications of motility more difficult compared to the control condition or the dsFoxP1 and dsFoxP2 conditions. Taken together, our findings are compatible with a role of FoxP transcription factors in the regulation of cell surface molecules that control the mode of cell-cell interactions and motility.

The observed phenotypes are also in agreement with the known functions of the FoxP transcription factors as homo-or hetero-dimers (Stroud et al., 2006; Villalobos et al., 2023). In this context, our failed attempts to rescue all the observed phenotypes after silencing the FoxP genes can be explained. The incomplete rescue of aberrant DRG phenotypes, even at high concentrations is consistent with the so-called classic transcription factor dosage paradox and the known requirement of FoxP transcription factors to function as homo-and heterodimers. High-dose rescue may disrupt normal FoxPx/FoxPy dimer formations as FoxP ratios are not restored and therefore more or fewer FoxP homo-versus heterodimers are formed. In turn, the perturbation of FoxP stoichiometry following knockdown may lead to the formation of nonfunctional or mis-specified dimers, thereby selectively impairing subsets of FoxP1-dependent transcriptional programs.

Still, in contrast to the DRG phenotype, for all three FoxP family members, low to intermediate levels of re-expression were sufficient to restore normal morphology of the sciatic nerve, indicating that FoxP activity is required in a cell-type specific manner. Notably, rescue efficiency varied both between FoxP paralogs and among DRG and the sciatic nerve, suggesting that individual cell types differ in their sensitivity to FoxP dosage and/or in their reliance on specific FoxP-dependent transcriptional programs. These findings indicate that FoxP transcription factors are not functionally interchangeable across all PNS development.

Together, these results demonstrate that FoxP proteins are essential regulators of PNS development, acting in a dose-sensitive and nerve-specific manner. Correct FoxP stoichiometry is required to support the full complement of peripheral nerve morphologies, and precise regulation of FoxP levels is critical for normal establishment of the peripheral circuitry. Our findings thereby provide a link that could explain why patients diagnosed with neurodevelopmental disorders often suffer from aberrant sensory perception that involve the PNS. This has been described for autism spectrum disorders, where beyond the core diagnostic features of autism, atypical behavioral responses to sensory information are observed in over 90% of autistic individuals (Tavassoli et al., 2014). Historically, in comparison to the research on the core symptoms of autism, little work has been done investigating the mechanisms behind the somatosensory abnormalities associated with ASD. However, sensory symptoms may not only precede but also be predictive of social-communication deficits and repetitive behaviors. For example, studies point out that tactile experiences during early childhood play a critical role in the development of normal social behavior and communication skills in both humans and rodents (Mammen et al., 2015). Individuals with autism spectrum disorder (ASD) often exhibit altered pain responsivity, which may be related to ASD-associated sensory differences or to atypical processing of nociceptive stimuli. These abnormalities are typically attributed to alterations in central sensory processing, particularly in the multimodal integration of sensory information. However, given the pervasiveness of sensory symptoms in ASD, the potential contribution of the peripheral sensory nervous system should not be overlooked. Consistent with this possibility, recent studies using skin biopsies have shown that approximately 53% of individuals with autism exhibit reduced intraepidermal nerve fiber density (Chien et al., 2020). This work is only a part of a broader trend in neuroscience to look beyond the brain to investigate how neuronal alterations outside the CNS may help to explain a host of the condition’s characteristic traits.

## Materials and methods

Double-stranded RNA (dsRNA), ISH probes, and plasmids dsRNAs were prepared by in vitro transcription from plasmids containing cDNA fragments (expressed sequence tags [ESTs] obtained from Source BioScience) as described previously (Pekarik et al., 2003). The ESTs used in this study are listed in Table 1. Chicken ESTs were linearized by digestion with the restriction enzymes NotI, EcoRI and EcoRV, and transcribed into DIG-labeled antisense and sense probes with T3 and T7 RNA polymerases, respectively (Mauti et al., 2006).

**Table 1.**
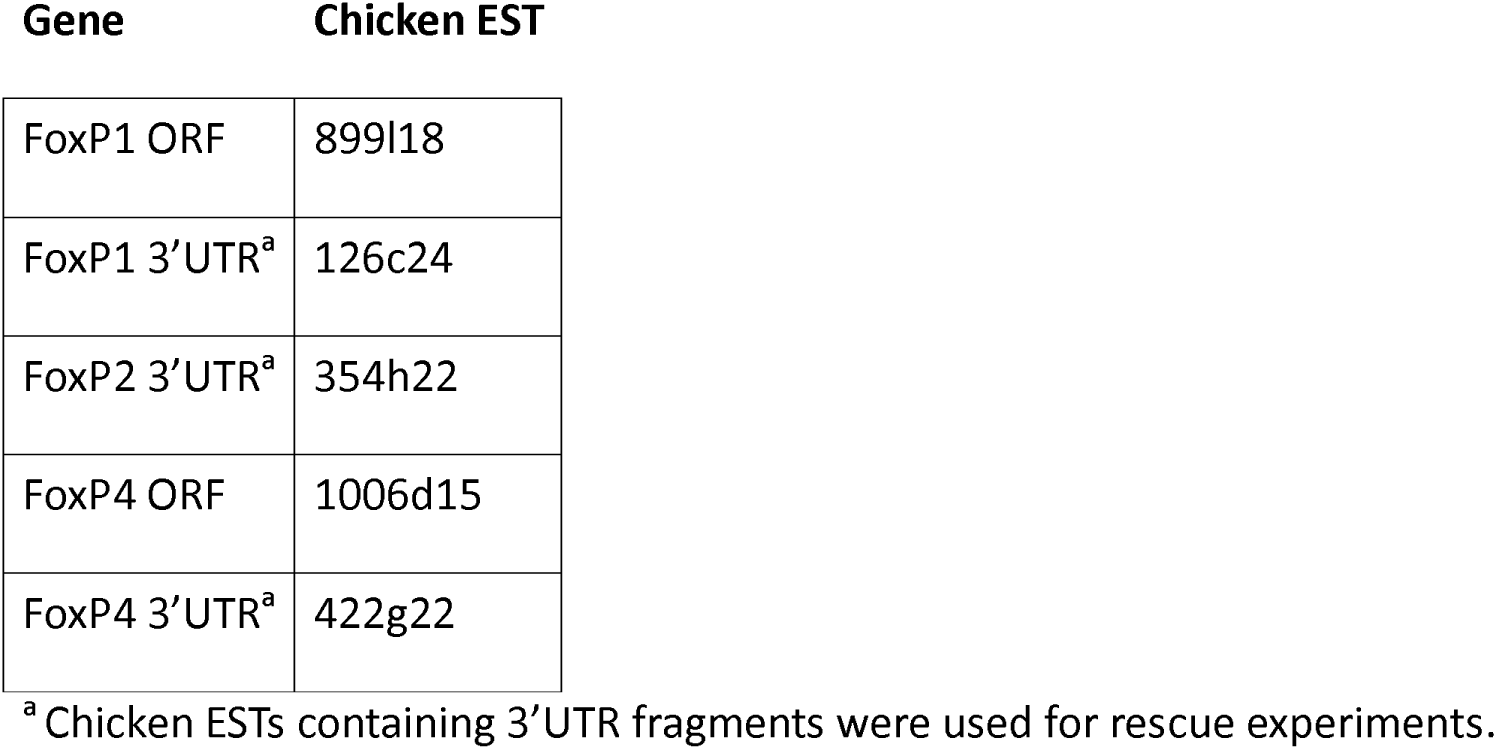
Chicken ESTs and cDNAs used to generate ISH probes and dsRNA Gene Chicken EST.

**Gene Chicken EST**
|  |  |
| --- | --- |
| FoxP1 ORF | 899l18 |
| FoxP1 3'UTR<sup>a</sup> | 126c24 |
| FoxP2 3'UTR<sup>a</sup> | 354h22 |
| FoxP4 ORF | 1006d15 |
| FoxP4 3'UTR<sup>a</sup> | 422g22 |

<sup>a</sup>Chicken ESTs containing 3'UTR fragments were used for rescue experiments.

For the rescue plasmids, we cloned the open reading frames of mouse FoxP1 (BioCat. 20794014-ABM), mouse FoxP2 (BioCat. 20797014-ABM), and mouse FoxP4 (BioCat. 20799015-ABM) into the respective vector containing the sequence for the 3xHA-(FoxP1), the 6xMyc-(FoxP2), or the 3xFLAG-tag (FoxP4). All constructs were verified by Sanger sequencing and successful expression of the tagged versions of FoxP transcription factors were tested in transverse sections of embryos electroporated with the rescue construct and the GFP-expression plasmids (Supplementary Figure 1).

### In ovo RNAi

All experiments involving animals were according to the guidelines of the Cantonal Veterinary Office Zurich. After 2 d of incubation at 39°C, chicken embryos were electroporated (as described in Table 2) after injection into the central canal of the spinal cord of either 20 ng/µl of plasmid containing a β-actin-driven GFP reporter alone (control), or together with 500 ng/µl of long dsRNA derived from either FoxP1, FoxP2, or FoxP4. For rescue experiments various concentrations of mFoxP1, mFoxP2, mFoxP4-containing constructs were co-injected with the mixes used for the downregulation as described in Table 2. Depending on the experiment, chicken embryos were sacrificed at various developmental stages (staged according to Hamburger and Hamilton, 1951) for whole-mount staining or sectioning.

**Table 2.** Concentrations and electroporation parameters of plasmids and dsRNA used in this study^a^.

**Name Concentration Electroporation**
| Name | Concentration | Electroporation |
| --- | --- | --- |
| dsFoxP1 | 500 ng/μl | 5 pulses, unilateral, 18 V<br>3 pulses, bilateral, 18 V <sup>b</sup> |
| dsFoxP2 | 500 ng/μl | 5 pulses, unilateral, 18 V<br>3 pulses, bilateral, 18 V <sup>b</sup> |
| dsFoxP4 | 500 ng/μl | 5 pulses, unilateral, 18 V<br>3 pulses, bilateral, 18 V <sup>b</sup> |
| β-actin::GFP | 20 ng/μl | 5 pulses, unilateral, 18 V<br>3 pulses, bilateral, 18 V <sup>b</sup> |
| β-actin::mFoxP1-3xHA | 50-250 ng/μl | 5 pulses, unilateral, 18 V |
| β-actin::mFoxP2-3xFLAG | 50-250 ng/μl | 5 pulses, unilateral, 18 V |

<sup>a</sup>All dsRNA and plasmid constructs were co-injected with 20 ng/ μl of β-actin::GFP. The interpulse interval was 1 second.
<sup>b</sup>For the analysis of knockdown efficiency by Western blots (WB) and for the live-imaging of delaminating NCCs from neural tube explants embryos were electroporated bilaterally, 1 s interpulse interval.

### In situ hybridization and immunohistochemistry

Chicken embryos were sacrificed at the desired stages (Hamburger and Hamilton, 1951). Dissected spinal cords were fixed in 4% of paraformaldehyde in PBS for 45 min, incubated overnight in 25% sucrose, and frozen in OCT. The tissue was cut into 25-µm-thick transverse cryosections. In situ hybridization (ISH) was performed as previously described (Mauti et al., 2006) with ISH probes used at a concentration of 1 ng/µl. All solutions were prepared with water treated with diethyl pyrocarbonate to protect RNA from degradation.

Immunohistochemistry was performed as previously described (Vaccaro et al., 2022). Sections were incubated in blocking solution (5% fetal calf serum in PBS with 0.25% Triton) for 1.5h prior to incubation overnight with the primary antibodies diluted in blocking solution. Primary and secondary antibodies used in this study are listed in Table 3.

**Table 3.**
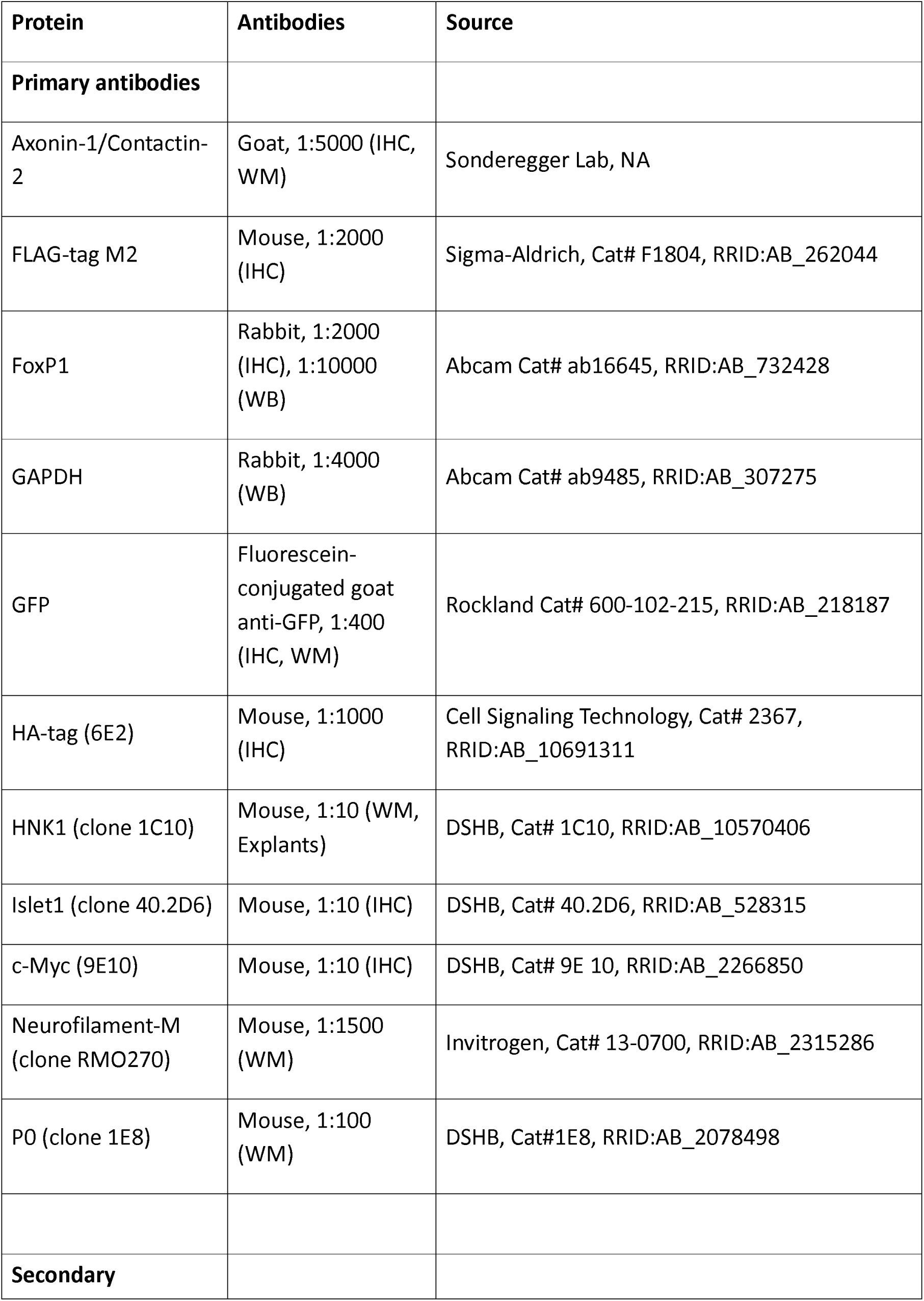
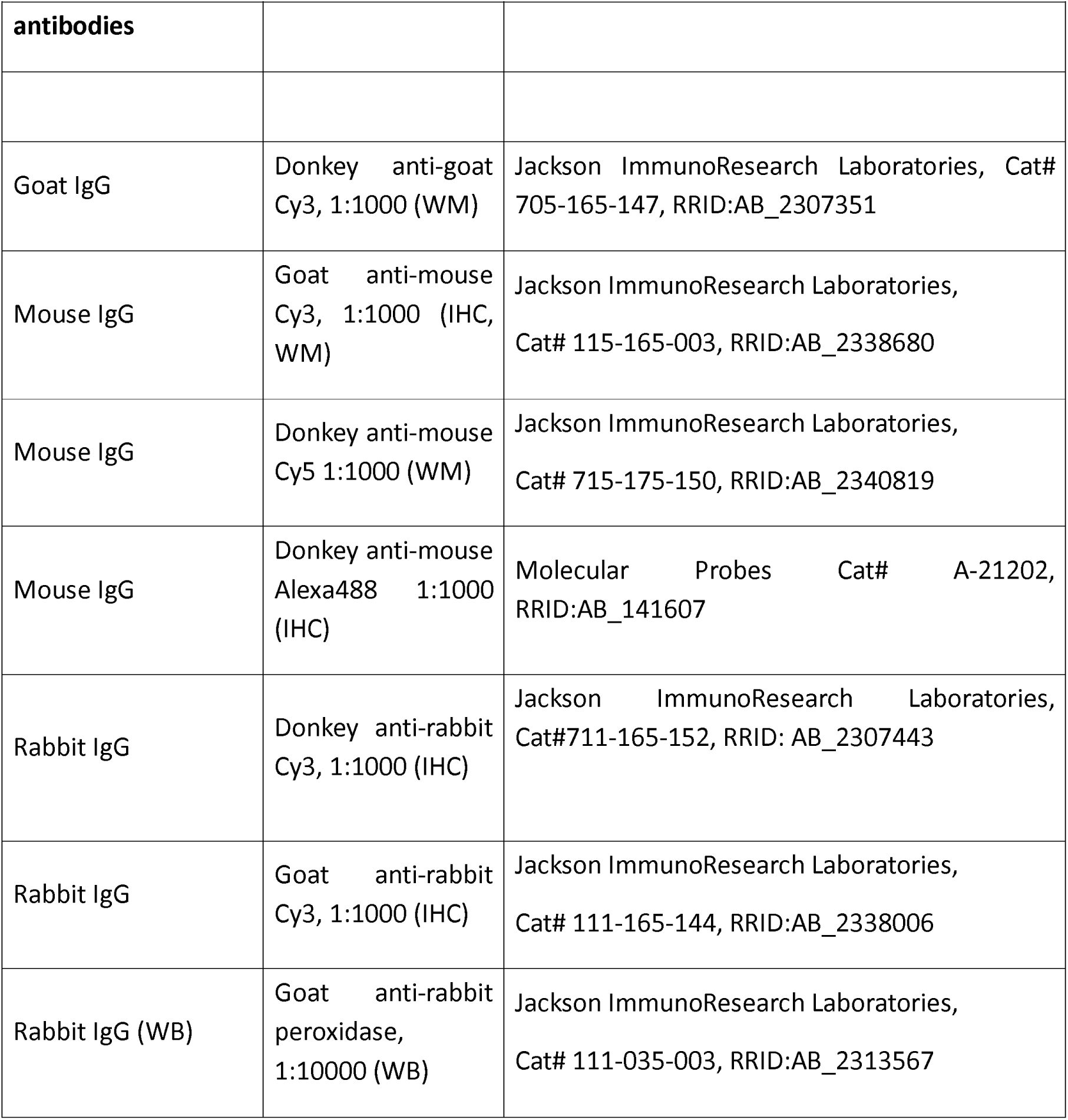

### Immunoblotting

Chicken embryos were injected and electroporated bilaterally at E3 (HH17-18) with 20 ng/µl of the GFP expression plasmid alone (control) or together with 500 ng/µl of dsFoxP1. After 3 d of incubation, embryos were sacrificed and spinal cords were collected in lysis buffer (125 mM NaCl, 10% glycerol, 1% NP40, 1 mM CaCl_2_, 1 mM MgCl_2_, 5 mM NaF, 1 mM Na_3_VO_4_, 10 mM β-glycerol phosphate, 1 tablet of protease inhibitors/10 ml in 20 mM Tris-Cl, pH 7.6) on ice. After trituration and centrifugation for 5 min at 4°C, supernatant and pellet were collected separately. After protein quantification, with a BCA assay, 50 µg of spinal cord lysates were loaded per lane onto an 8% SDS gel. The protein bands were transferred to nitrocellulose membranes. Membranes were washed in TBST before using the ECL Western Blotting Detection Reagent (GE Healthcare). The chemiluminescence signal was detected using the Amersham Imager 600 (GE Healthcare). Resulting bands were quantified through ImageJ. Antibodies used in these experiments are listed in Table 3.

### Neural tube explants

HH11-HH13 chicken embryos were injected with 50 ng/μl β-actin::H2B-mGreenLantern (produced from addgene plasmid #164464) and 600 ng/μl crestin::tdTomato-F before bilateral electroporation with 3 pulses at 18 V each side. Ibidi 8-well plates (Ibidi, #80806-90) were coated with bovine Fibronectin (20 μg/ml in PBS, pH 7.4). HH17 embryos were dissected in cold PBS, pH 7.4. Embryos were incubated in Dispase II (1 mg/ml in sterile PBS) at room temperature for 10 min, followed by PBS washes. Tungsten wires were used to remove the neural tube from the mesenchyme between leg and wing levels. Then, neural tubes were sliced into pieces of a length of 2 somites with scalpel blades, and lumbar level explants were transferred to the wells filled with 250 μl live-imaging medium (Neurobasal medium, 4 mg/ml Albumax [Gibco], 1 mM sodium pyruvate [Sigma], 100 units/ml Penicillin, B27 [Invitrogen, 17504044], and 2 mM L-Glutamine). Neural tube explants were plated dorsal side down and allowed to adhere to the wells for 2h at 37 °C before live imaging was started.

### Microscopy

Images of spinal cord cryostat sections were acquired using an Olympus BX61 upright microscope. Imaging was performed with a 10× air objective (UPLFL PH 10×/0.30) or a 20× air objective (UPLXAPO 20x/0.8), using Olympus CellSens Dimension software.

Visualization of HNK1-positive neural crest cells (NCCs) in HH18 whole-mount embryos was performed using an Olympus IX83 inverted microscope equipped with a spinning disk unit (CSU-X1, 10,000 rpm; Yokogawa), with 10× and 20× air objectives.

Time-lapse imaging of migrating neural crest cells was conducted using the same Olympus IX83 spinning disk microscope. Cultured neural tube explants were maintained at 37 °C in a humidified atmosphere containing 5% CO₂ and 95% air using a PeCon CellVivo chamber (PeCon). Explants were allowed to equilibrate for 2 h prior to imaging. Images were acquired every 7 min over a 20-h period, using a 10× air objective and an Orca-Flash 4.0 camera (Hamamatsu), controlled by Olympus CellSens Dimension software.

For imaging of BABB-or ECi-cleared whole-mount embryos, a mesoSPIM light-sheet microscope was used (mesoSPIM V6, camera Hamamatsu Orca Lightning; Voigt et al., 2019; Vladimirov et al., 2024). Images were acquired at 2x magnification (Mitutoyo M Plan Apo 2x/0.055 objective, effective pixel size 2.75 µm/px, z-step size 5 µm), or at 5× magnification (Mitutoyo M Plan Apo 5x/0.14, effective pixel size 1.1 µm/px, z-step size 5 µm).

The samples were pinned to a slab of agarose which was immersed in a cuvette (Portmann Instruments) filled with index matching medium (BABB or ECi) for imaging. In the detection path we used fluorescence filter QuadLine Rejectionband ZET 405 / 488 / 561 / 640 (AHF# F57-405SG).

### Staining and tissue clearing of whole-mount embryos

After fixation, embryos were transferred to a 24-well plate and stored overnight in PBS at 4 °C. The following day, embryos were permeabilized in 1% Triton X-100 in PBS for 1 h, followed by incubation in 20 mM lysine in 0.1 M sodium phosphate buffer (pH 7.3) for 1 h at room temperature. Embryos were then rinsed five times in PBS (10 min per wash). To reduce nonspecific binding, samples were incubated in blocking buffer (10% fetal calf serum in PBS) for 2 h, after which primary antibodies diluted in blocking buffer were applied for 48 h at 4 °C. Embryos were subsequently washed in PBS with five solution changes (2 h each), followed by an additional overnight wash at 4 °C. After a further 2 h incubation in blocking buffer, embryos were incubated with the appropriate secondary antibodies overnight. Samples were then washed five times in PBS and incubated overnight at 4 °C. Embryos were dehydrated either through a methanol gradient for BABB clearing or through an ethanol gradient for ECi clearing (25%, 50%, 75% in H₂O, followed by two changes of 100%; 1 h per step; ethanol pH 9–9.5). Finally, embryos were cleared by incubation in a 1:2 mixture of benzyl alcohol and benzyl benzoate (BABB) or in ethyl cinnamate, overnight at 4 °C (BABB) or at room temperature (ECi).

### Whole-mount embryos quantifications and statistical analysis

For downregulation and rescue experiments, cleared embryos were scored blinded to the experimental treatment. For each nerve, the assessment was performed visually based on well-defined stereotyped morphological criteria of each nerve. For NCC polarity assessment, HH18 embryos were assessed in a blinded manner based on well-described NCC delamination pattern characteristics as normal (0) or aberrant (1) polarity. Binary data were stored in a CSV file and imported into Python using Pandas. Phenotypic data was subsequently analyzed in Python using pandas, matplotlib, and seaborn. For each experimental condition and nerve, the proportion of embryos with aberrant phenotypes was calculated as the mean of the binary scoring values. These proportions were visualized using stacked bar plots displaying the fraction of normal and aberrant embryos per condition. Since the data consist of binary categorical outcomes and some groups contained relatively small sample sizes Fisher’s exact test was selected. Binary values (0 = normal, 1 = aberrant) were visualized as a heatmap using a two-color scale. Pairwise statistical comparisons between experimental groups were performed using the two-sided Fisher’s exact test implemented in SciPy. For each comparison, a 2 × 2 contingency table was constructed using the counts of aberrant and normal embryos in the two groups. Significance thresholds were defined as ns, when p ≥ 0.05; *p < 0.05; **p < 0.01; ***p < 0.001; ****p < 0.0001. Resulting p values were corrected for multiple testing using the Benjamini–Hochberg false discovery rate (FDR) procedure. Volcano plot for the downregulation experiment was generated with seaborn and matplotlib, plotting effect size against the negative log10-transformed FDR-adjusted p value. A horizontal dashed line indicated the significance threshold at FDR = 0.05.

### Neural crest cells tracking and cell trajectory reconstruction

Cell migration was analyzed by manually tracking individual cells over time using time-lapse microscopy images in Fiji/Image J Manual tracking plugin. For each explant, 15-20 individual cells located at the migrating front were randomly selected and traced. For the assessment of the 1^st^ delaminating wave, the tracing was done for 6h of recordings (14-20h). Cell tracking was performed by recording the centroid position of each cell at every time point. Tracking data were exported as.csv files containing cell ID, frame/time, coordinates. Representative tracks were visualized using Python (pandas and matplotlib) by reconstructing trajectories from sequential coordinates. Cell trajectories were reconstructed, optionally rotated for orientation consistency, and visualized as static track plots and animated GIFs (Movies 4-7). Animations were rendered using Matplotlib FuncAnimation and exported with PillowWriter.

## Statistical analysis

Downregulation efficiency: For each embryo, multiple measurements of the electroporated/control ratio were obtained and averaged to generate one value per embryo. Embryo means were treated as biological replicates. Data are presented as mean ± SD. Comparisons were performed between GFP_HH25 and dsFoxP1, GFP_HH29 and dsFoxP2, and GFP_HH18 and dsFoxP4. Normality was assessed using the Shapiro–Wilk test (n ≥ 3). If both groups were normally distributed, an unpaired two-tailed Welch’s t-test was used; otherwise, a two-sided Mann–Whitney U test was applied. Significance thresholds were defined as ns, p ≥ 0.05; *p < 0.05; **p < 0.01. Downregulation efficiency was calculated as (mean GFP − mean dsFoxP) / mean GFP) × 100. Statistical analyses were performed in Python using SciPy, and plots were generated using Matplotlib.

Neural crest cells analysis: Track-summary data were imported into Python and analyzed using pandas, NumPy, SciPy, Matplotlib, and Seaborn. Conditions were ordered as Control, dsFoxP1, dsFoxP2, and dsFoxP4. For each track, four migration parameters were examined: absolute displacement, total distance, straightness index, and average speed. Data distributions were visualized as boxplots with overlaid jittered single-track measurements, colored by explant, and explant-level means were indicated as larger markers. Control was compared with each dsFoxP condition using Welch’s two-sample t-tests. Significance was annotated on plots as ns (P ≥ 0.05), * (P < 0.05), ** (P < 0.01), or *** (P < 0.001).

The authors have no conflict of interest

## Acknowledgements

We would like to thank Dr. Beat Kunz and Tiziana Flego for excellent technical assistance. The project was supported by a fellowship Swiss Government Excellence scholarship (ESKAS scholarship) and by the University Research Priority Program ‘Adaptive Circuits in Development and Learning’ (URPP AdaBD) of the University of Zurich.

## Supplementary Data

**Supplementary Figure 1.**
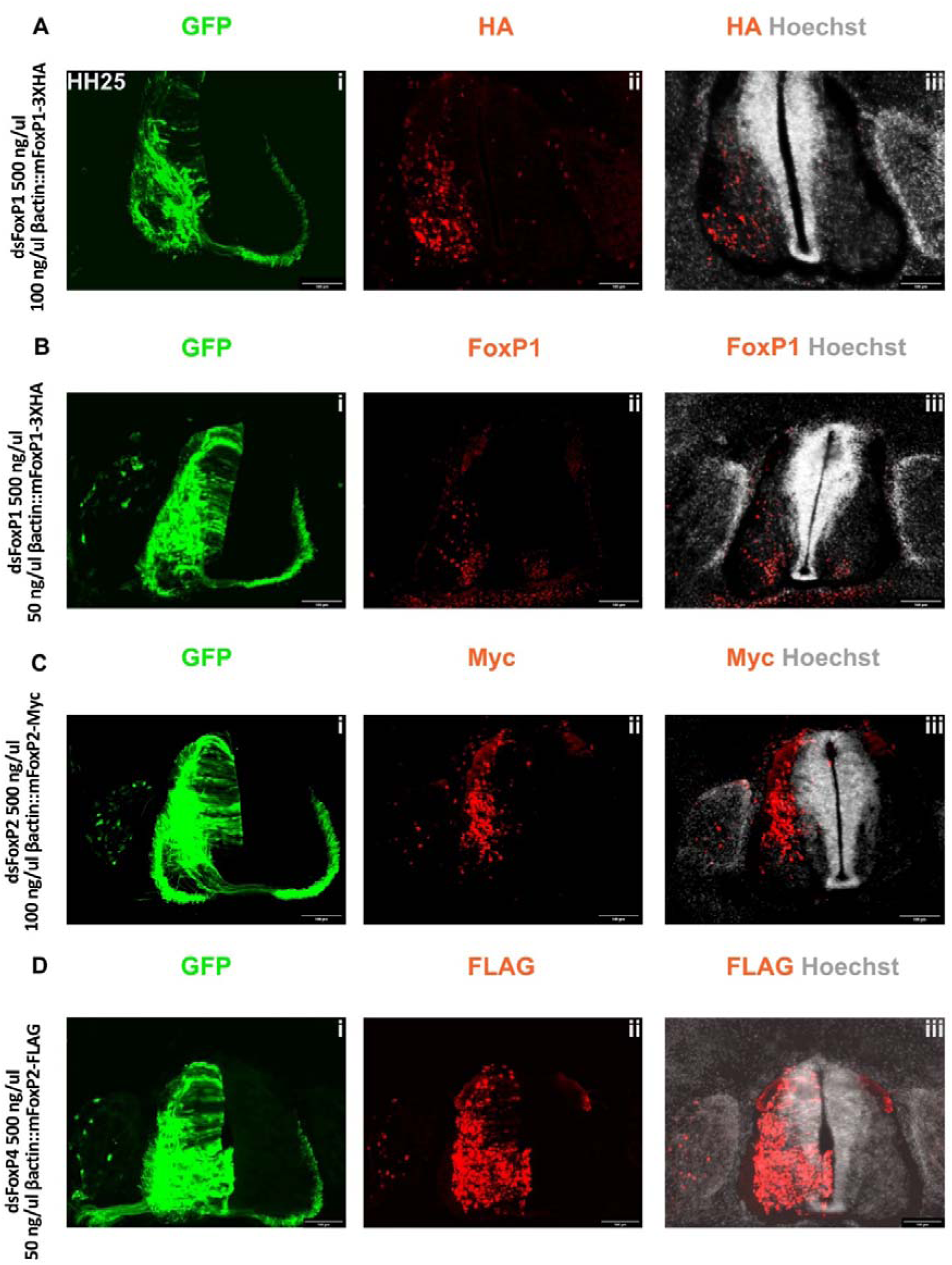
Detection of tagged rescue constructs in cryosections. Transverse spinal cord sections were stained with antibodies against the tags, as indicated. In all embryos, a GFP expression plasmid (β-actin::GFP; 20 ng/µl) was included in the injection mix to verify successful electroporation. Sections were counterstained with Hoechst. (A) The expression of mFoxP1 (100 ng/µl) in an embryo co-injected and electroporated with dsFoxP1 (500 ng/µl) reveals successful expression of mFoxP1-HA (ii). (B) With an antibody recognizing FoxP1, we could directly verify successful expression of FoxP at the protein level. Note that the levels and the distribution between the left (electroporated) and the non-electroporated side differ in (B). Successful expression of the mFoxP2-myc (C) and the mFoxP4-FLAG (D) constructs was demonstrated by staining for the myc-tag and the FLAG-tag, respectively. Scale bars: 100 µm.

## Movies

**Movie 1. The PNS of a whole-mount GFP-expressing control embryo at HH25 stained with anti-Neurofilament antibodies.** After clearing with BABB, the embryo was imaged with a mesoSPIM light-sheet microscope to visualize the neural circuits in the hindlimb and DRG.

**Movie 2. Embryos lacking FoxP transcription factors exhibit aberrant PNS development.** Compared to the GFP-expressing control embryo shown in Movie 1, the embryo lacking FoxP4 exhibits aberrant hindlimb innervation. For example, the ventral crural nerve was defasciculated to various degrees on both sides.

**Movie 3. The migration pattern of delaminating neural crest cells is impaired after silencing FoxP genes.** Live imaging of neural crest cells migrating from explants on a fibronectin-coated dishes reveals differences in migration behavior. Using manual tracking, we followed the movements of selected NCCs. In control explants, NCC migration is characterized by cell-cell contacts via transient processes followed by changes in migration direction after contact. Downregulation of FoxP1 caused NCCs to travel shorter distances. NCCs lacking FoxP2 had fewer cell-cell contacts. They had the same displacement as controls, but slower speed and straighter trajectories. Cells lacking FoxP4 clustered during migration and stayed in contact with other cells by forming tight cell-cell contacts involving large surface areas. This clustering behavior was not interfering with migration as the displacement and the total distance were not different from controls, despite the fact that the cells moved as a cluster. Scale bar: 100 µm. Fiji, manual tracking.

**Movie 4. Tracks of NCCs from control explants. Using manual tracking, we followed the movements of selected NCCs and reconstructed their pathways using custom Python coding.** Cell trajectories were reconstructed, optionally rotated for orientation consistency, and visualized as animated GIFs showing progressive migration over time. Each cell has an individual track color. Behavior is described in Movie 3.

**Movie 5. Tracks of NCCs from explants taken from dsFoxP1 embryos.** Using manual tracing, we followed the movements of selected NCCs and reconstructed their pathways using custom Python coding. Cell trajectories were reconstructed, optionally rotated for orientation consistency, and visualized as animated GIFs showing progressive migration over time. Each cell has an individual track color. Behavior is described in Movie 3.

**Movie 6. Tracks of NCCs from explants taken from dsFoxP2 embryos. Using manual tracing, we followed the movements of selected NCCs and reconstructed their pathways using custom Python coding.** Cell trajectories were reconstructed, optionally rotated for orientation consistency, and visualized as animated GIFs showing progressive migration over time. Each cell has an individual track color. Behavior is described in Movie 3.

**Movie 7. Tracks of NCCs from explants taken from dsFoxP4 embryos. Using manual tracing, we followed the movements of selected NCCs and reconstructed their pathways using custom Python coding.** Cell trajectories were reconstructed, optionally rotated for orientation consistency, and visualized as animated GIFs showing progressive migration over time. Each cell has an individual track color. Behavior is described in Movie 3.

